# From model to field: predictive design of microbial communities for resilient crop performance

**DOI:** 10.1101/2025.10.21.683643

**Authors:** Shinichi Yamazaki, Masaru Nakayasu, Keiko Kanai, Rie Mizuno, Rumi Kaida, Sachiko Masuda, Arisa Shibata, Ken Shirasu, Atsushi J. Nagano, Yoshiharu Fujii, Akifumi Sugiyama, Yuichi Aoki

## Abstract

Synthetic microbial communities (SynComs) represent a promising approach to enhance crop growth and stress resilience through microbiome engineering. However, the systematic design and field validation of SynComs remain limited. Here, we present a predictive framework for SynCom optimization, integrating plant phenotyping, microbial genomics, and machine learning. Using tomato as a model, we tested over 800 SynCom–temperature combinations consisting of root endophytic bacteria and rhizosphere metabolites. An Elastic Net regression model trained on plant biomass data accurately predicted the performance of unseen SynComs, with prediction accuracy plateauing at ∼5% (301/6144) of all possible SynCom–temperature combinations. Incorporating genomic features significantly improved model performance, whereas microbiome compositional data alone were not informative. We applied the model to design novel SynComs, which were tested in both laboratory and field conditions using a commercial tomato cultivar. The model-informed SynCom enhanced plant growth in field trials and improved heat stress tolerance under controlled laboratory conditions. Multi-omics analyses and feature importance metrics identified specific microbial taxa, including Sphingobium sp., whose enrichment was linked to host plant metabolite (e.g., tomatine) and stress-responsive gene expression. Our results demonstrate a scalable strategy for the predictive design of beneficial microbiomes to improve resilient crop performance under real-world conditions.

## Introduction

The plant rhizosphere is a major site of plant–soil–microbe interactions and constitutes a unique chemical environment that contains a unique mix of various metabolites, organic compounds, and minerals. Root-associated microorganisms influence plant nutrient acquisition, pathogen resistance, and abiotic stress sensitivity ^1^. These effects mean that plant rhizosphere microbiota is an important factor regulating the growth of host plants. Using microbes to improve plant nutrient use efficiency, alleviate environmental stress, and suppress plant pathogens has been proposed as a strategy to reduce the use of chemical fertilizers and pesticides in agriculture, thereby enhancing sustainable food production ^2–4^.

The rhizosphere contains a microbial community that differs from that present in bulk soil, and this is shaped by various metabolites secreted by plant roots. Root exudates contain over 10% of the carbon that plants fix via photosynthesis ^5,6^, and their primary metabolites, including sugars, organic acids, and amino acids, are essential carbon and nitrogen sources for soil microorganisms ^7,8^. In recent years, researchers have found that specialized (secondary) metabolites are involved in the formation of various plant-specific rhizosphere microbiota. For example, tomatine, a saponin possessing antifungal, antibacterial, and insecticidal activity that is produced by tomato (*Solanum lycopersicum*), has been found to be involved in establishing and maintaining the rhizosphere microflora, where it increases the abundance of a specific bacterial group (e.g., genus *Sphingobium* ^9^). Recently, our team revealed that another isolate from tomato roots, *Sphingobium* sp. RC1, participates in a tomatine degradation pathway, which suggests that it may be involved in interactions specific to tomato ^10^. Other compounds, including the soybean isoflavonoid and maize flavanone ^11,12^, the sulfur-containing indole alkaloid camalexin produced by *Arabidopsis thaliana* ^13^, coumarin secreted by *A. thaliana* ^14^, and the benzoxazinoids produced by maize ^15,16^, are now thought to be involved in the shaping of the rhizosphere microbial community and the regulation of plant growth promotion by microbes. Furthermore, recent studies have shown that metabolite and microbial profiles within the rhizosphere are also altered by abiotic stress ^17,18^, suggesting that plants may modulate rhizosphere microbiota by varying root-secreted metabolite production to facilitate acclimation to various environmental changes. These findings highlight the potential to engineer rhizosphere microbial communities that support plant resilience under abiotic stress conditions.

Many plant growth-promoting (PGP) microorganisms have also been isolated from plant rhizospheres ^19^. Known PGP traits include phytohormone production, increased nutrient availability via biological nitrogen fixation and solubilization of insoluble minerals, and biological control of pathogens via antibiotic and siderophore production ^20–22^. Experimental work has shown that inoculating a single strain with a specific PGP trait is a simple method that is effective in specific environments; however, the effect may not always be consistent ^23,24^. This unpredictability can occur due to oversimplified models of the effects of plant-microbe associations, and can result in inoculated microbes being quickly outcompeted by species in the indigenous microbiome ^23,25^. In addition, the soil microbiota creates a diverse and complex network of interactions that can make it difficult to understand the detailed mechanisms at work. One approach to resolving this issue involves the use of synthetic microbial communities (SynCom). SynCom is an emerging technique involving the coculturing of multiple microbial strains to mimic the structure and function of environmental microbial communities ^26,27^. This approach allows us to determine the function of complex microbial communities and their biological interactions by isolating them as simple systems. For example, Castrillo *et al*. used SynCom to demonstrate the association among phosphate starvation response, immune response, and root microbiome in Arabidopsis ^28^, while Niu *et al.* elucidated the role-specific dynamics of representative bacteria found in the maize root microbiome by using simplified SynCom as a model ecosystem ^29^. In addition, SynCom has been found to produce superior pathogen suppression effects than single strain inoculation, indicating that constituent bacteria may exert synergistic effects ^30^. However, when designing a SynCom using different microbial compositions, it may be costly or impossible to experimentally verify all possible combinations and thereby identify the best SynCom. In such a case, it may be more helpful to make use of computational approaches such as machine learning to optimize modeling and prediction. In recent years, some studies have applied machine learning techniques to omics datasets (including the microbiome, transcriptome, metabolome, and genome, among others) to predict soil health indices ^31^ and to identify genetic and/or metabolic biomarkers associated with human disease ^32,33^.

In this study, we aimed to develop a model-based approach for optimizing SynComs to promote plant growth and improve crop resilience under environmental stress conditions. We designed a diverse SynCom composed of tomato root endophytic bacteria, including *Sphingobium* sp. RC1, in combination with rhizosphere metabolites. Tomatine was added alongside the SynCom, based on previous findings that microbial metabolic capacity can influence plant-microbiota interactions in the rhizosphere ^34–37^. Next, we constructed machine learning models by combining various SynCom compositions with genomic and microbiome information to predict plant growth and design optimal SynComs. Feature importance of constructed models and multi-omics analysis of SynCom-inoculated plants were used to identify key microbes associated with plant growth promotion. Finally, we investigated the molecular mechanisms underlying these interactions and their role in supporting crop performance under stress conditions (Fig. 1).

**Figure 1.**
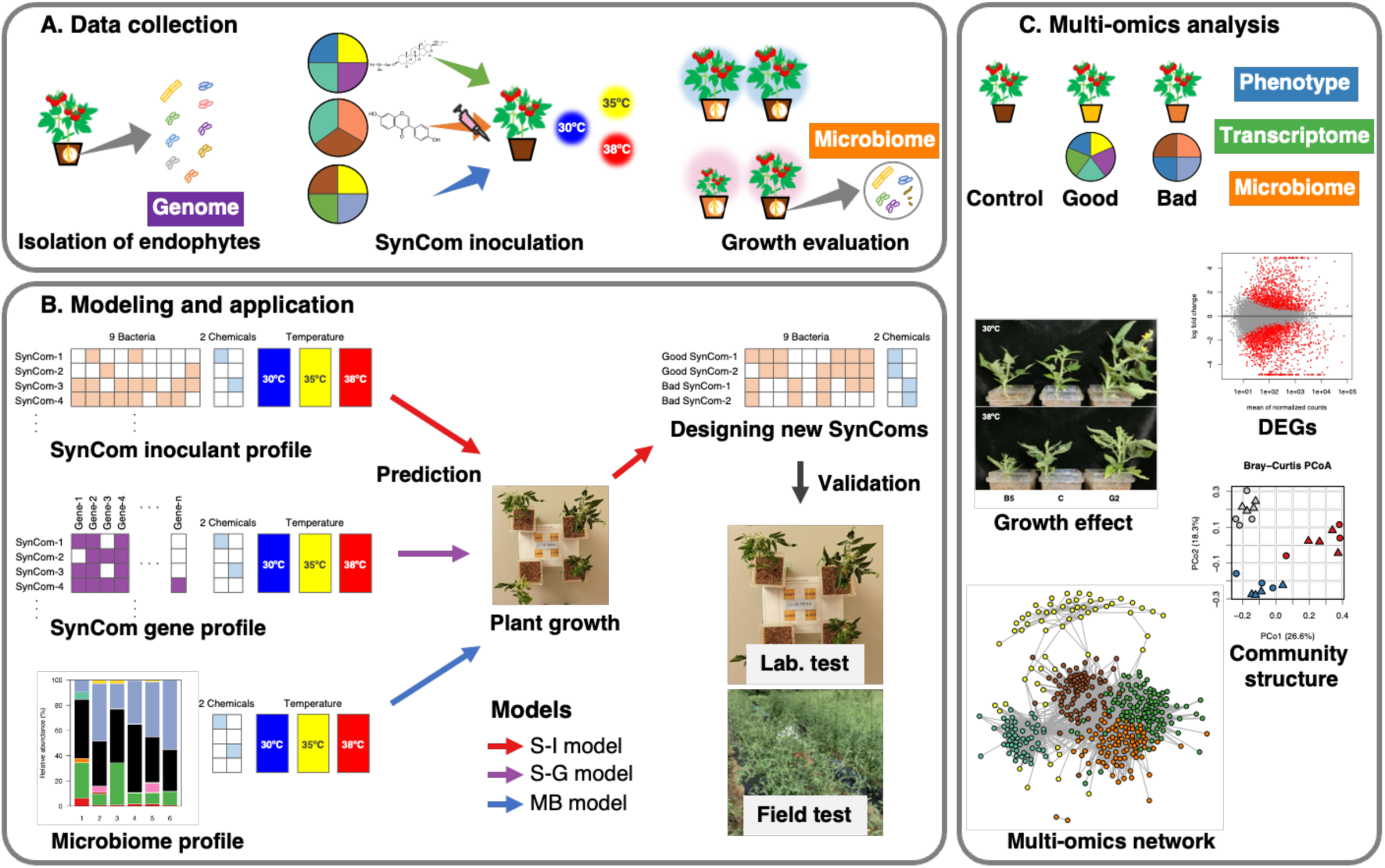
Overview of this study. **A.** Data collection. Endophytic bacteria were isolated from tomato roots, then various SynCom compositions were prepared and inoculated into tomato plants to examine their effects on plant growth. We also performed genome analysis of isolated endophytic bacteria and microbiome analysis of SynCom-inoculated tomato plants. **B.** Modeling and application. Based on data collected, we created three models to predict plant growth. The first model used the SynCom bacterial composition as an explanatory variable, the second model used the SynCom genetic composition, and the third model used the actual microbiome composition. New SynComs were then designed by constructing predictive models; these were then validated in laboratory and field cultivation experiments. **C.** Multi-omics analysis of SynCom-inoculated tomato plants. Phenotype, transcriptome, and microbiome data of inoculated tomato plants were obtained for newly-designed “Good” and “Bad” SynCom to validate their predicted effects on plant growth, changes in gene expression, and effects on microbial community structure. We also estimated the important genetic and microbial factors related to tomato growth using integrated multi-omics network analyses.

## Results

### Data collection of plant growth, root microbiome, and microbial genome

We first isolated 125 bacterial strains from the tomato root endophytic community. Taxonomic classification of these strains based on partial 16S rRNA gene sequences revealed that they belonged to four phyla (i.e., Actinobacteriota, Bacteroidota, Firmicutes, and Proteobacteria), representing 17 families and 26 genera (Supplemental Table S1). According to microbiome data produced by a previous study from our team ^9^ (DRA011415), most of these taxa were enriched in the tomato rhizosphere and root endosphere. We therefore selected nine phylogenetically diverse strains, including *Sphingobium* sp. RC1, from all strains that could be precultured from a stock (Fig. 2A).

**Figure 2.**
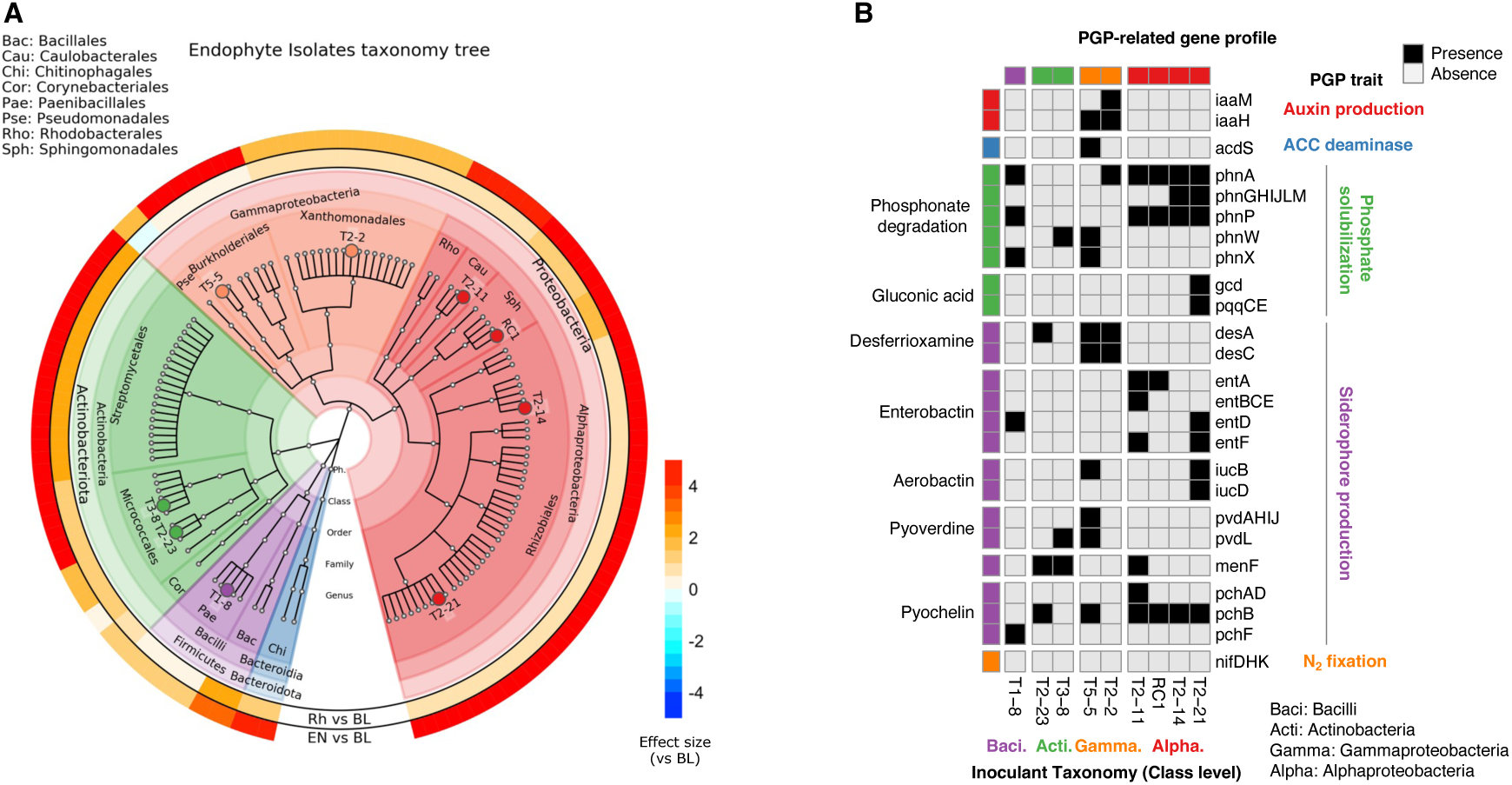
Phylogenetic and genetic characteristics of bacterial isolates from the tomato root endophytic community. **A.** Taxonomic tree of bacterial isolates from the tomato root endophytic community. Each isolate was assigned to a genus-level taxon by BLAST query of partial 16S rRNA sequences. The taxonomic hierarchical tree was then drawn using GraPhlAn. Strain names of selected SynCom members are indicated in the tree. Circular heatmaps outside the tree show differences in relative abundance of family-level taxa between bulk soil and rhizosphere or bulk soil and root endosphere, as indicated in previous data. The effect sizes estimated by ALDEx2 are shown as differences. Red indicates enrichment in the rhizosphere or root endosphere. **B.** Profiles of plant growth-promoting (PGP) trait-related genes carried by SynCom members. Genes for PGP trait-related pathways were determined by querying the MetaCyc and KEGG databases, and their presence in each genome was estimated.

We performed whole-genome sequencing analysis on all nine isolates. Overall we obtained nine high-quality circular genomes. We then attempted to identify these species by comparing sample genomes to known bacterial genomes. Two strains (T1-8 and T2-2) appeared as hits when querying the genomes of known species with an average nucleotide identity of more than 95%. For this analysis, four strains (T2-21, T2-23, T3-8, and T5-5) were marked as coming from the genomes of unspecified species, but other three strains (T2-11, T2-14, and RC1) did not hit any known genome and were considered to be new species (Supplemental Table S2). Gene prediction and functional annotation of the genome were then performed to characterize the genetic composition of each isolate. We compared the possession of genes related to PGP traits such as auxin production, the suppression of ethylene accumulation, phosphate solubilization, siderophore production, and nitrogen fixation ^20–22^ among the genomes of the nine isolates discussed here (Fig. 2B). With respect to plant hormones, T2-2 possessed the auxin biosynthetic genes *iaaM* and *iaaH*, while T5-5 possessed the ACC deaminase gene *acdS*, which can degrade ethylene precursor and suppress ethylene accumulation ^38^. As for phosphate solubilization, all the isolates except T2-23 possessed any of the *phn* genes that are involved in the degradation of phosphonate ^39^. In particular, two Rhizobiales isolates, T2-14 and T2-21, were found to possess more *phn* genes than any others. T2-21 is also linked to a gluconic acid biosynthetic pathway that is involved in phosphate solubilization. Regarding siderophore production, all nine isolates possessed at least one of the genes involved in known siderophore biosynthesis. Moreover, four isolates (i.e., T2-2, T5-5, T2-11, and T2-21) possessed multiple genes within these biosynthetic pathways. Finally, none of the isolates possessed the *nif* gene required for biological nitrogen fixation. Taken together, these results suggest that all nine isolates may be potential plant growth-promoting bacteria.

The resulting SynCom, which consisted of nine isolates, was then inoculated into tomato seedlings. These seedlings were grown under high temperature stress in a growth chamber, and the effect of the SynCom on plant growth was determined by evaluating root and shoot fresh weight. We also varied the SynCom composition, including various combinations of the nine isolates, for example excluding 1 or 2 isolates, or including only 1 or 2. SynCom inoculation was combined with addition of the plant metabolites tomatine or daidzein, and resulted in 2048 theoretical SynCom compositions. Next, we conducted 30 independent growing trials at 30°C (optimal), 35°C, and 38°C (high temperature), thus obtaining growth data for a total of 834 tomato plants accompanied by 218 SynCom combinations (Supplemental Table S3). In this series of trials, bacterial contamination was suspected in some of the microbial stocks used for SynCom preparation, so some of the strains used for the SynCom preparation were renewed for an 11th trial. In addition, since we observed some seasonal changes in growth data even under a controlled environment, a control condition (i.e., noninoculated, 30°C) was added from the 18th trial to facilitate batch-to-batch correction (Supplemental Fig. S1). Overall, we obtained growth data for 301 plants of 102 different SynCom compositions to be used for subsequent modeling.

We performed microbiome analysis via 16S rRNA amplicon sequencing of the endophytes of all tomato roots after taking root growth measurements. The relative abundance of amplicon sequence variants (ASVs) that corresponded to each of the nine inoculants was found to increase following inoculation, confirming that all inoculants had colonized their respective tomato roots (Supplemental Fig. S2). We also observed that plant metabolites exerted an effect, with simultaneous inoculation with tomatine significantly increasing the relative abundance of RC1 (Fig. 3A). In contrast no such change was observed for the other eight bacterial strains or during simultaneous inoculation with daidzein (Supplemental Fig. S3 and S4). Next, diversity analysis was performed to examine the effects of temperature and plant metabolite on the endophytic community. Overall, high temperature did not affect species richness and evenness, but the addition of plant metabolites tended to increase both diversity indices (Fig. 3B). Principal coordinate analysis (PCoA) based on weighted UniFrac distance showed a significant effect of temperature on endophytic community (PERMANOVA: p < 0.001, R^2^ = 0.021), but we did not observe a clear separation by temperature alone (Supplemental Fig. S5). Taken together, these results suggest that the effect of high temperature on the endophytic community is not a determinant factor and can be controlled by SynCom inoculation.

**Figure 3.**
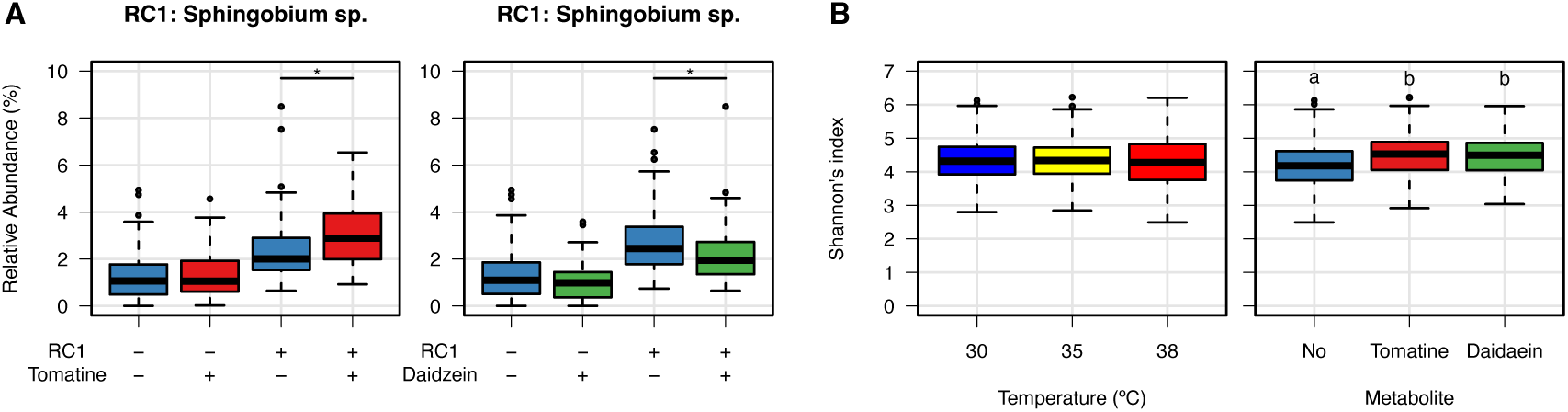
Microbiome analysis of SynCom-inoculated tomato roots used for model construction. **A.** Effect of metabolite addition on RC1 colonization on roots. Asterisk (*) indicates a significant difference in group means (p < 0.05, Wilcoxon rank-sum test). **B.** Effect of growing temperature and metabolite addition on the alpha diversity of root-associated bacterial communities. Different letters indicate significant differences between their mean values (p < 0.05, Wilcoxon rank-sum test with Bonferroni correction).

### Modeling and application

Based on the data collected, we constructed three different models to predict tomato growth. Specifically, we used SynCom inoculant composition (SynCom-inoculant model; S-I model), SynCom gene composition (SynCom-Genome model; S-G model), and microbial composition of the roots (Microbiome model; MB model) as explanatory variables, respectively. The algorithm we chose was an Elastic Net regression model, and this was then used to evaluate and compare the performance of the three models above. First, for the S-I model explanatory variables were divided into three categories: microbes, chemicals (i.e., tomatine and daidzein), and environment (temperature). Each of their respective contributions to overall plant growth was examined based on model performance. We found that environmental factors alone had the largest effect on plant growth (E: Mean R^2^ = 0.402), while microbial and chemical factors alone each had very small effects (M: Mean R^2^ = 0.046; C: Mean R^2^ = 0.010; Fig. 4A). However, the model containing all three factors as well as the interaction of two variables (MCEX) showed the highest performance (Mean R^2^ = 0.510). This suggested that microbial and chemical factors can both affect plant growth, and so we used all of these as explanatory variables in subsequent models.

**Figure 4.**
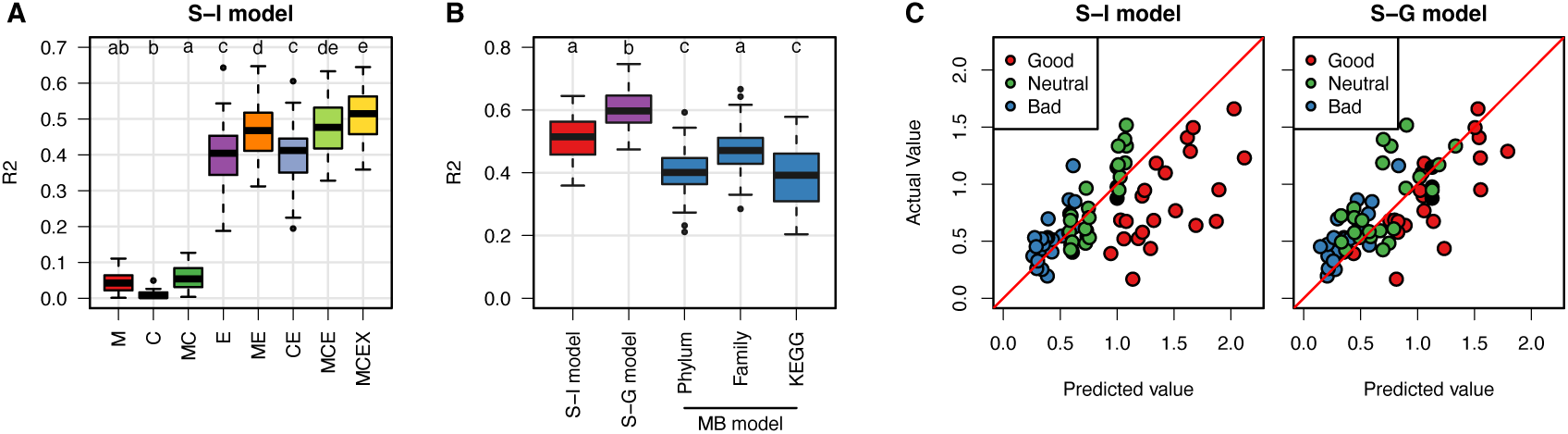
Summary of model performance. **A.** Effects of microbial, chemical, and environmental factors on plant growth prediction using the S-I model. M, microbial factor; C, chemical factor; E, environmental factor; X, interaction of two variables. **B.** Comparison of the predictive performance of the S-I, S-G, and MB models. Splitting of training and test data (i.e., in a 7:3 ratio) was randomly repeated 50 times to evaluate average model performance. Different letters indicate significant differences between group means (p < 0.05, Welch’s t-test with Bonferroni correction). **C.** Verification of the versatility of the S-I and S-G models. Correlation between predicted and measured values of Good (top 5%), Neutral (median) and Bad (bottom 5%) SynComs designed using the S-I model.

The S-G model using the SynCom genome information showed significantly better prediction performance (Mean R^2^ = 0.600) than the S-I model, and showed the highest performance among all models developed in this study (Fig. 4B). This result means that the gene composition of SynCom and especially the presence of specific functional genes strongly influence tomato growth. In contrast, using root microbiome data did not improve the predictive performance of the MB model (Mean R^2^ = 0.467) relative to the S-I model. Instead, models using phylum-level profiles and inferred metabolic pathway profiles showed significantly lower performance (Mean R^2^ = 0.407; Mean R^2^ = 0.384). These results suggest that microbiome information sampled after cultivation is insufficiently able to explain the biomass of host plants and that the composition of SynCom inoculants and/or genomic data is crucial.

We also that, in addition to the Elastic Net algorithm used in most models, we also tested Lasso and Random Forest algorithms. However, in all three models (i.e., S-I, S-G, and MB) the Elastic Net algorithm consistently showed the best performance (Supplemental Fig. S6). We also examined the effect of sample size by stepwise subsampling of training data. In doing so we found that the Elastic Net model was mostly saturated in predictive performance using the full size of the current dataset, indicating that the sample size of our dataset was sufficient for modeling (Supplemental Fig. S7).

To verify the versatility of constructed models, we designed new SynComs based on the S-I model from the top, middle, and bottom predicted values. We then acquired new growth data and evaluated the predictive performance of the models (Fig. 4C). Overall, a significant correlation was observed between measured and predicted values for the S-I model (p = 8.704e-09, r = 0.619), and an even higher correlation was obtained for the S-G model (p = 2.815e-13, r = 0.735). These results suggest that it may be useful to design a new SynCom using the machine learning model, and that more effective SynCom design and modification can be achieved by using genetic information from constituent bacteria.

Based on the feature importance of the S-I model, we identified specific candidate microbe-microbe and microbe-plant metabolite interactions that may promote plant growth. The top 15 variables deemed most important for positive association with tomato growth were microbial interactions, either with each other or with metabolites (Fig. 5A,B). Of these, the simultaneous inoculation of two isolates—e.g., T2-2 and RC1 or T3-8 and T5-5—was found to significantly improve tomato growth when cultivated at either optimal (30°C) or high (35°C) temperature (Fig. 5B,C). In addition, the simultaneous addition of RC1 and tomatine microbe-metabolite pair was found to significantly improve heat stress tolerance (Fig. 5C). Taken together, these results show that such interactions have a synergistic effect on plant growth promotion.

**Figure 5.**
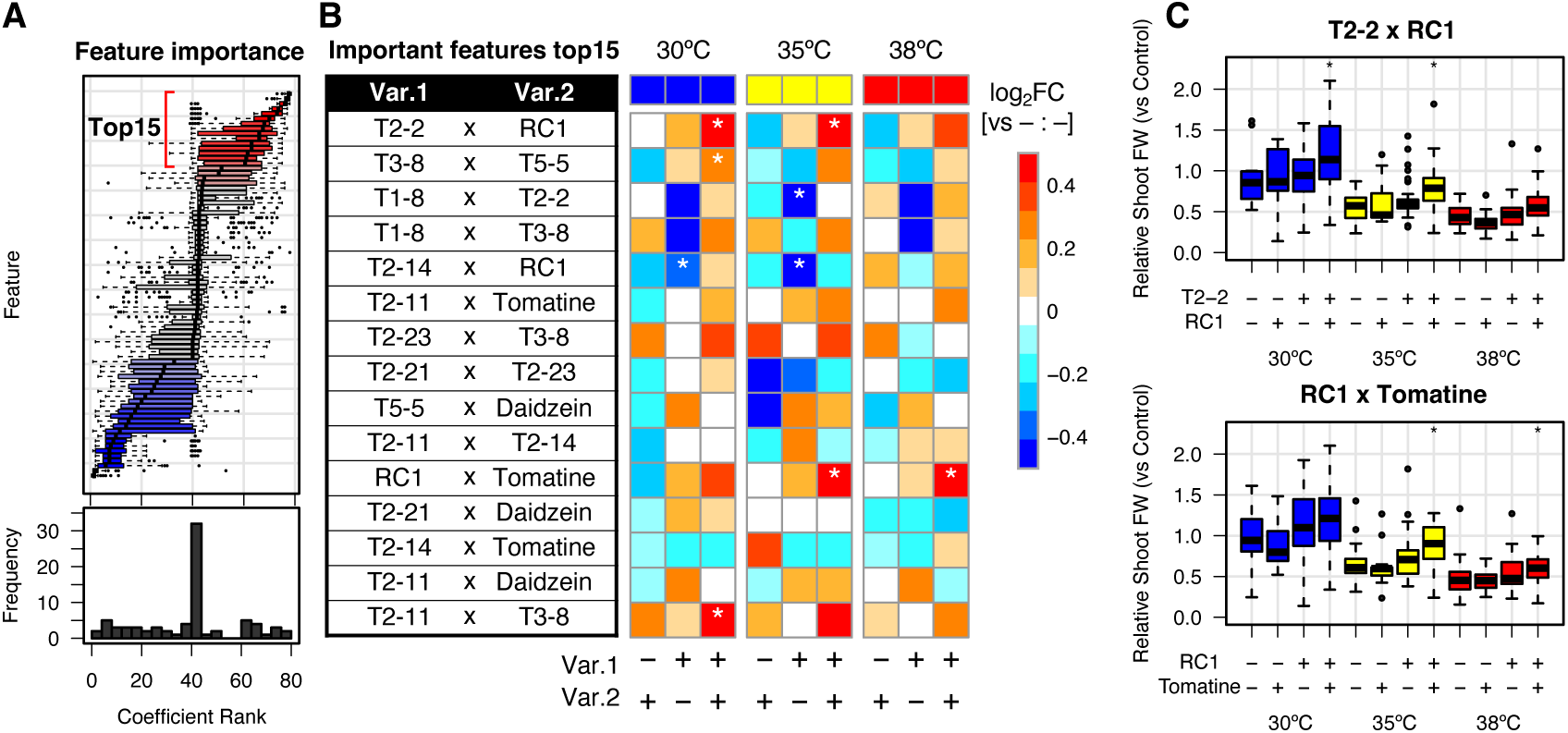
Feature importance of the SynCom-Inoculant model. **A.** Distribution of regression coefficients for the S-I model. Here, coefficients are ranked in ascending order, and the distribution and median of rankings over 50 iterations are shown in box plots and a histogram. **B.** Top 15 positive important variables for tomato growth and their effect sizes. They are all two variable interactions, and the heatmap shows the effect sizes of single or double variables on plant growth. Values indicate logarithms of median fold change relative to the minus:minus group. Asterisks (*) indicate significant difference relative to the minus:minus group (p < 0.05, rank-based Dunnett test). Var, variable; FC, fold change. **C.** Plant growth enhancement by microbe-microbe (T2-2 and RC1 strains) and microbe-chemical (RC1 and tomatine) interactions.

Although we also attempted to functionally characterize the important genes of the S-G model by performing GO enrichment analysis, this was discarded since only about 10% of all genes were annotated using GO terms, which was not sufficiently informative.

### Multi-omics analysis of tomato plants inoculated with SynCom

The top five SynComs designed based on the S-I model, hereafter termed Good SynCom G1–G5, were evaluated again using a field cultivation experiment to determine their effects on plant growth. Most of the Good SynComs tended to increase plant biomass. In particular, SynCom G2 showed a statistically significant growth-promoting effect relative to non-inoculated plants (Fig. 6A). Overall, SynCom G2 consisted of six bacterial strains (i.e., T2-2, T2-11, T2-23, T3-8, T5-5, and RC1) combined with tomatine. We subsequently performed a detailed multi-omics analysis of SynCom G2 using laboratory experiments. These results were then compared to non-inoculated and Bad SynCom B5 preparations, and inoculated tomatoes were grown under optimal and high temperature conditions to generate plant biomass, leaf and root transcriptome, and root microbiome data. SynCom B5 consisted of tomatine and five bacterial strains (i.e., T1-8, T2-14, T2-21, T2-23, and T5-5) and was among the bottom five SynComs generated based on the S-I model. Overall, SynCom G2 inoculation was associated with a significant improvement in plant tolerance to heat stress, whereas SynCom B5 inoculation resulted in no differences relative to non-inoculated plants (Fig. 6B–D).

**Figure 6.**
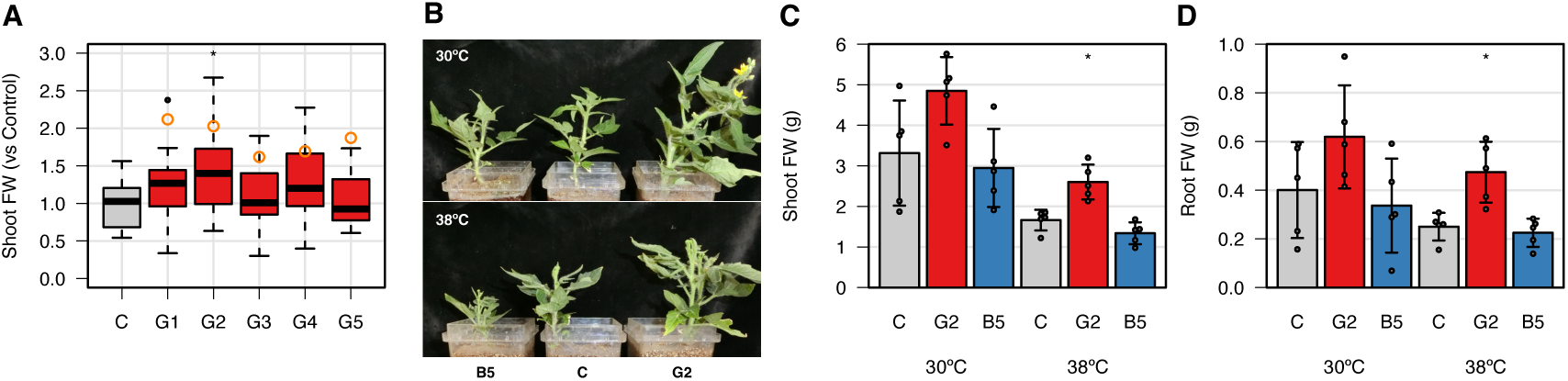
Validation of the predicted beneficial effect of Good SynCom on tomato plant growth. **A.** Field growth of Good SynCom G1-G5-inoculated tomato plants. Orange empty circles represent predicted values for the S-I model. The field was divided into four blocks, including four plots per treatment and four plants per plot (n = 16). Asterisk (*) indicates a significant difference relative to the uninoculated control (p < 0.05, Dunnett test). **B.** Representative growth of SynCom G2- and B5-inoculated tomato plants in the laboratory experiment. **C, D.** Shoot and root biomass of SynCom G2- and B5-inoculated tomato plants in the laboratory experiment. SynCom-inoculated tomato plants were grown under optimum (30°C) and high temperature (38°C) conditions (n = 5). Asterisks (*) indicate a significant difference relative to an uninoculated control (p < 0.05, Dunnett test).

In transcriptome analysis, principal component analysis and clustering analysis showed that tomato gene expression was clearly different based on tissue and temperature (Supplemental Figs. S8 and S9). Moreover, the effect of SynCom inoculation was remarkable in the roots exposed to heat stress conditions (Supplemental Fig. S8). Next, we conducted a differential gene expression analysis between SynCom-inoculated and non-inoculated tomatoes. We found that the number of differentially expressed genes (DEGs) was highest in roots inoculated with SynCom G2 under heat stress conditions and was the lowest in roots inoculated with SynCom B5 at the same temperature (Supplemental Fig. S10). Next, DEGs were clustered based on their coexpression patterns and functionally characterized by gene ontology (GO) enrichment analysis. These results showed that, for root tissue subjected to heat stress conditions, G2 SynCom inoculation upregulated the expression of genes involved in “response to heat” and “protein folding” (Fig. 7). For example, we found significant up-regulation of heat shock transcription factors (HSFs) and heat shock proteins (HSPs), two important regulators of heat stress response and protein quality, respectively (Fig. 8). Such changes in gene expression may have improved the heat tolerance of the tomato plants. Moreover, we also found that SynCom B5 inoculation increased the expression of genes involved in “response to biotic stimulus” in leaves (Fig. 7), thereby suggesting that this SynCom exerts an unfavorable effect on tomato plants.

**Figure 7.**
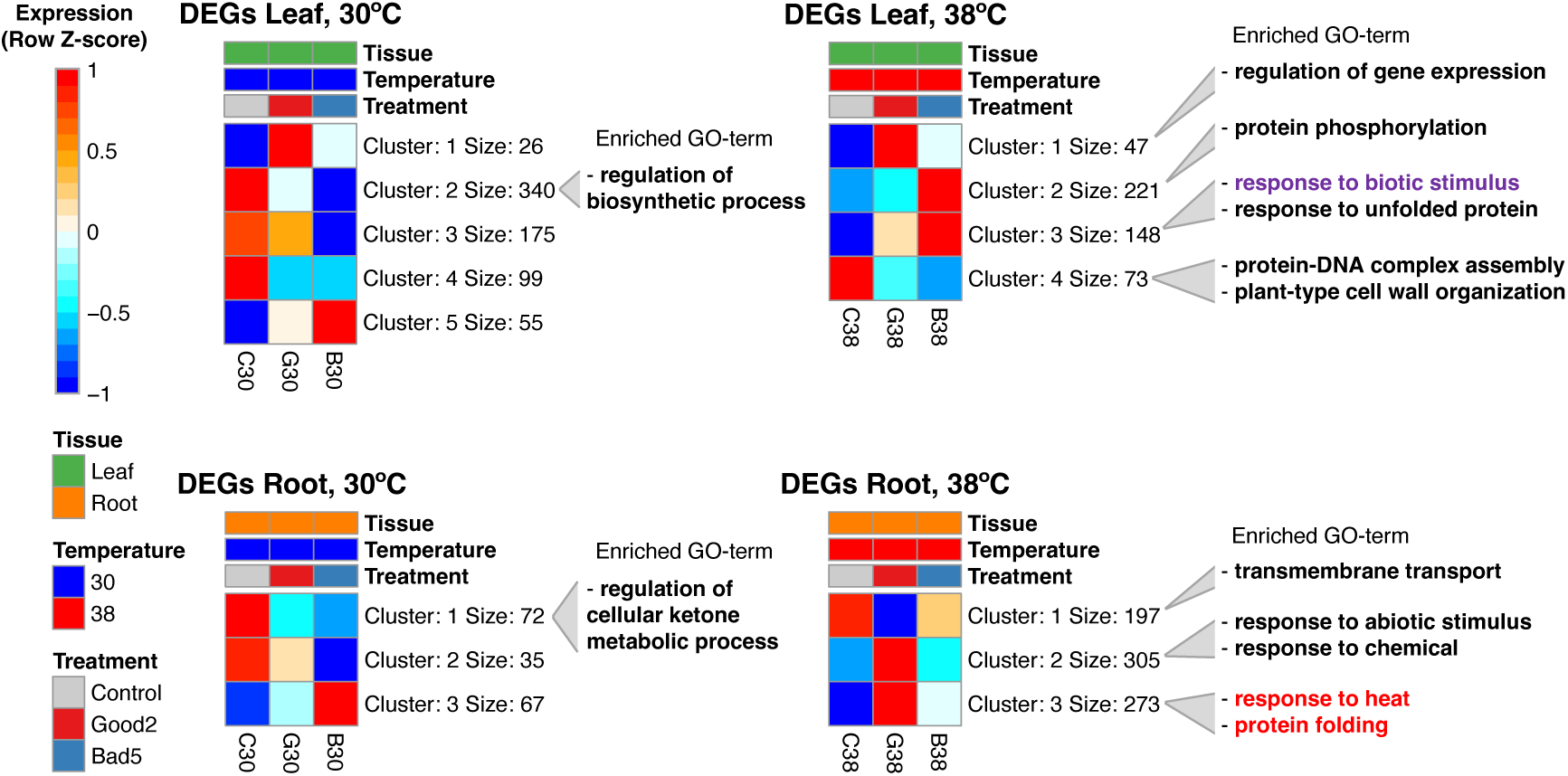
Expression pattern clustering and GO-term enrichment of differentially expressed genes (DEGs) responsive to SynCom inoculation. DEGs related to SynCom G2 and B5 inoculation were identified for each tissue and temperature (FDR < 0.01, DESeq2). Next, these DEGs were clustered using the k-means method. The optimal k was determined based on the Bayesian information criterion. Representative gene expression patterns within each cluster were revealed by the heatmap. GO enrichment analysis was performed for each gene cluster (FDR < 0.05, hypergeometric test).

**Figure 8.**
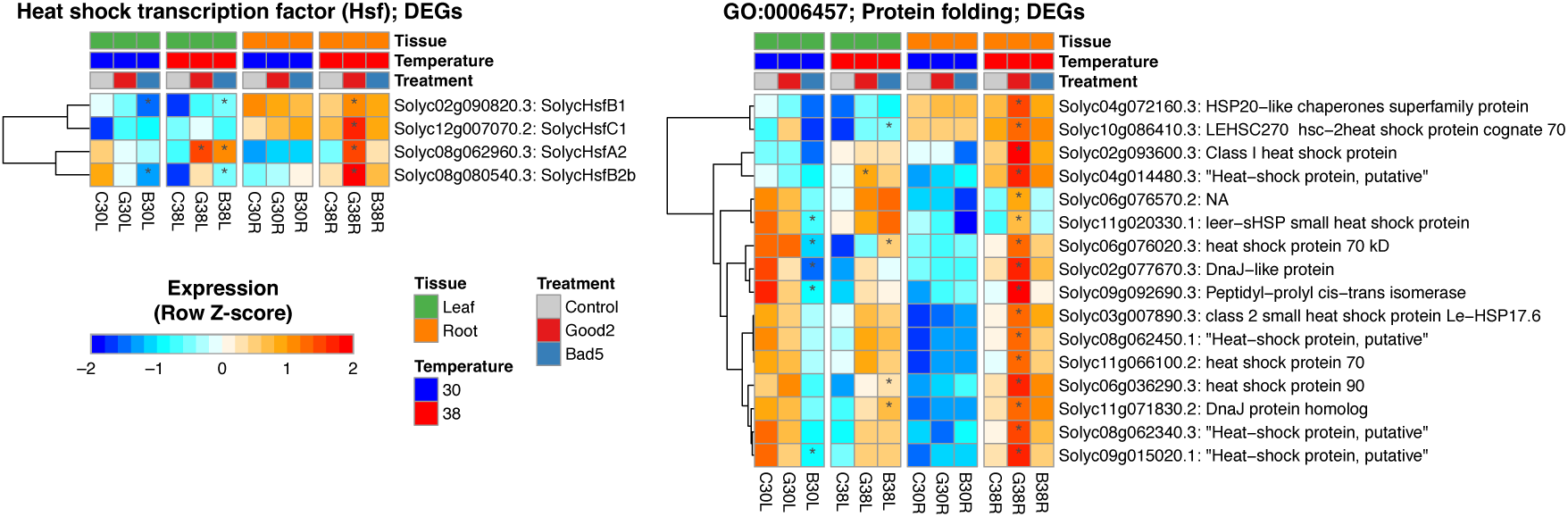
Expression profiles of HSF and HSP genes. Among all DEGs identified following SynCom inoculation, those annotated with the gene symbol “HSF” and the GO-term “protein folding” were extracted and their expression patterns are shown in the heatmap. Asterisks (*) indicate significant differences relative to an uninoculated control (FDR < 0.01, DESeq2).

In the microbiome analysis, a principal coordinates analysis (PCoA) based on Bray-Curtis dissimilarity distance and PERMANOVA revealed that the endophytic community of tomato roots was significantly affected by SynCom inoculation (p < 0.001, R^2^ = 0.410; Fig. 9A). In contrast, temperature showed a limited effect (p = 0.082, R^2^ = 0.051). Moreover, the alpha diversity of the endophytic community was decreased by SynCom G2 inoculation (Fig. 9B). In general, the composition of bacterial taxa on the genus level differed greatly among treatments, and SynCom members increased in abundance following inoculation (Supplemental Fig. S11). Especially in SynCom G2-inoculated tomatoes, *Pseudoxanthomonas* (same genus as T2-2) was the most dominant, accounting for 30% of the total (Fig. 9C). Differential abundance analysis using ALDEx2 showed significant increases in *Achromobacter* (same as T5-5), Micrococcaceae (same as T2-23) and *Pseudoxanthomonas* in SynCom G2 relative to the control (Fig. 9D). For SynCom B5, we observed significant increases in *Phyllobacterium* (same as T2-21) and *Achromobacter*. Of these, *Achromobacter* (T5-5) was twofold higher in SynCom G2-inoculated tomatoes than SynCom B5 inoculation. In addition, *Achromobacter* was the dominant genus (i.e., ranking in the top 15) only in the SynCom G2-inoculated tomatoes even though it was a common constituent in both SynCom G2 and B5.

**Figure 9.**
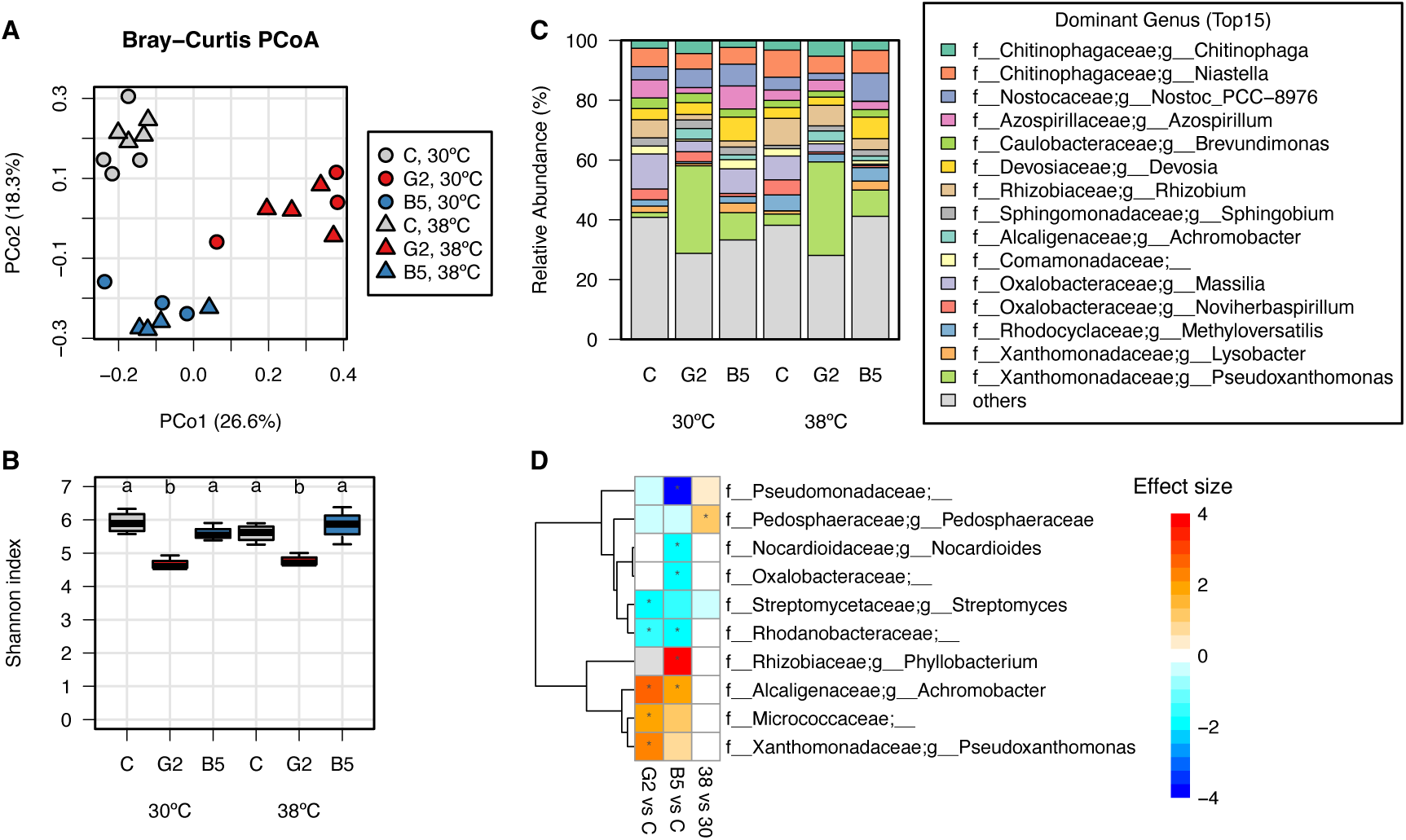
Root-associated microbial community structure following SynCom inoculation. **A.** Principal coordinate analysis (PCoA) plot of Bray–Curtis distance among samples. **B.** Shannon index representing the alpha diversity of the microbial community. Different letters indicate a significant difference between SynCom group means (p < 0.05, Welch’s t-test with Bonferroni correction; n = 3 for G2 and B5 at 30°C, n = 4 for the others). **C.** Relative abundance of the dominant top 15 taxa within the community resolved to the genus level. Values indicate the means of 3-4 biological replicates. **D.** Differentially abundant taxa on the genus level by SynCom inoculation or high temperature status. An ALDEx2 generalized linear model (aldex.glm) was used to test the effect of each factor, and effect sizes are shown by the heatmap. Asterisks (*) indicate significant differences relative to the control (FDR < 0.1, ALDEx2).

Finally, integrated omics analyses were performed using sparse canonical correlation analysis (sCCA) and co-occurrence network analysis. We extracted relevant components among different omics data for plant biomass, leaf and root transcriptome, and root microbiomes by sCCA. Next, co-occurrence network analysis using WGCNA was used to analyze the relationship between pairs of components and to examine the characteristic co-occurrence modules associated with plant biomass. Overall, sCCA was able to extract components of 4,938 leaf genes, 7,089 root genes, and 176 root bacterial taxa (genus level). Network analysis allowed the classification of the 12,205 components into 29 modules based on co-occurrence patterns. When considering the specific module associated with plant biomass (i.e., the fresh weight of shoots and roots), the network analysis identified four genes in leaves, 85 genes in roots, and one bacterial taxon, *Sphingobium*. Although we observed no significant enrichment of specific GO terms in this module, the gene Solyc05g053080.2, which encodes endomembrane-type CA-ATPase 4, was the hub and was found to be associated with *Sphingobium* and plant biomass (Fig. 10A). In addition, the relative abundance of *Sphingobium* and the gene expression of Solyc05g053080.2 were both strongly correlated with plant biomass (Fig. 10B). Taken together, these results indicate that *Sphingobium* spp. in the root microbiota are closely related to tomato growth and that specific plant genes may be involved in this relationship.

**Figure 10.**
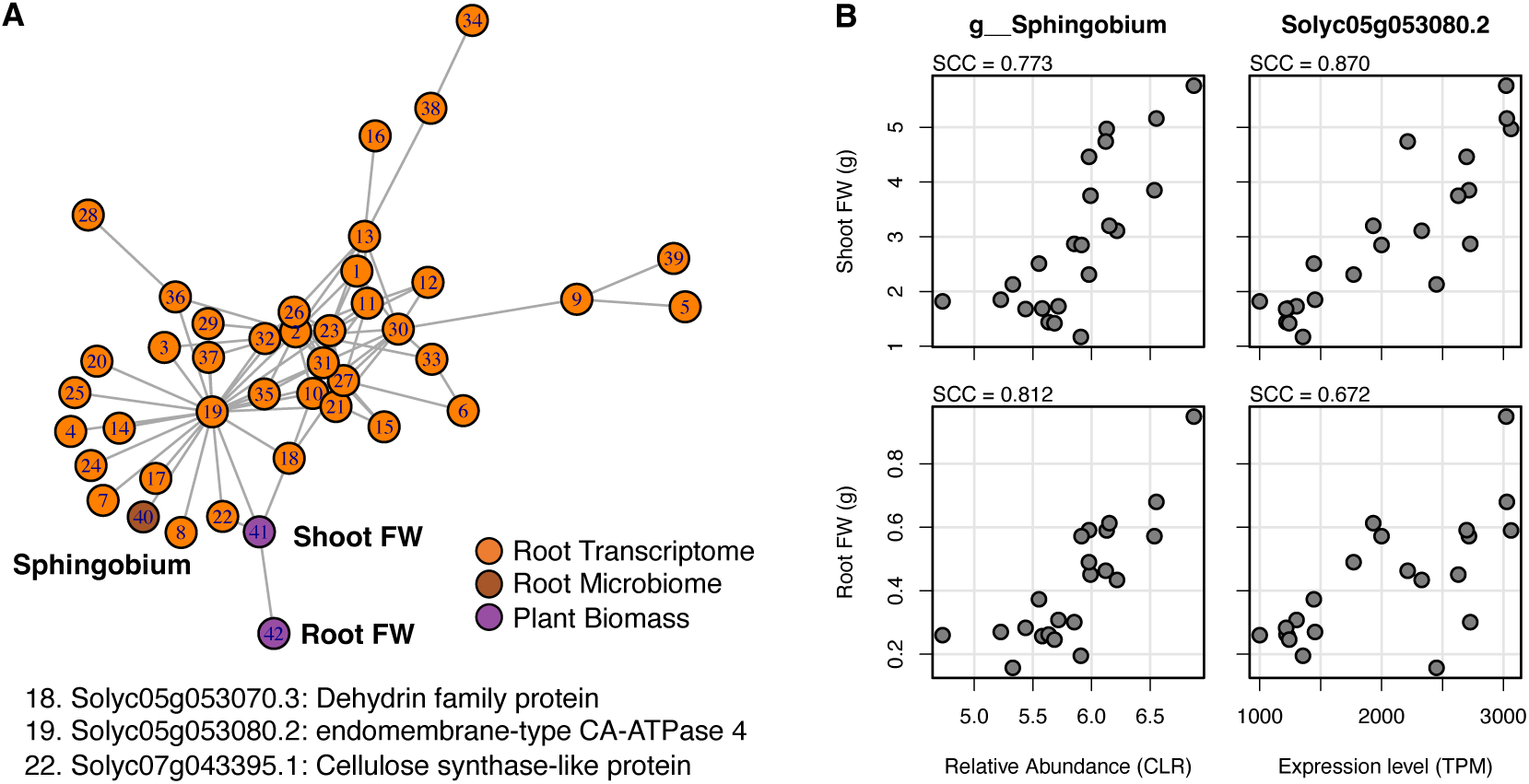
Plant biomass-associated modules in a multi-omics co-occurrence network. **A.** Sub-network of modules related to plant biomass, created using WGCNA. Nodes represent tomato genes, bacterial taxa on the genus level, and plant phenotypic traits. Edges were drawn based on the topological overlap matrix with a threshold of 0.08. **B.** Correlations between plant biomass and relative abundance of *Sphingobium* sp. as well as between plant biomass and the expression levels of the hub gene Solyc05g053080. SCC, Spearman correlation coefficient; FW, fresh weight.

## Discussion

To increase sustainability of crop production in an era of climate change, it is important to maximize crop production potential by making better use of soilborne microorganisms and metabolites produced by plants. One approach to achieve this is to mimic microbial communities using SynCom techniques ^26^. In this study, we used machine learning to optimize SynCom design, and followed this process from model construction to experimental validation (Fig. 1).

Specifically, we constructed models to predict plant growth based on SynCom structural information and achieved high prediction performance. These results show that it is feasible to design an arbitrary SynCom based on machine learning models. Here, we used relatively simple data and models (e.g., binary data describing inoculum presence/absence and linear models). We built the model to represent 50% of the plant biomass (Mean R^2^ = 0.510) from real data for about 5% of all SynCom combinations. Moreover, performance improvement was found to be almost saturated in the linear model (Supplemental Fig. S7), and we therefore conclude that there was little benefit in further increasing the sample size. This model may serve as a useful example for future studies applying similar approaches.

The SynCom-Genome model, which incorporates information regarding SynCom gene composition, showed the highest predictive performance in this study (Fig. 4B). This suggests that SynCom genetic composition, or the metabolic functions within the community, can strongly influence plant growth and abiotic stress tolerance. Although the metabolic activities of the community remain poorly understood, it has recently been reported that the exchange of extracellular metabolites, or metabolic interactions, can affect the community phenotype, e.g., drug resistance ^40^. In addition, recent studies have used mutant libraries to identify genetic requirements for the establishment of microbial interactions ^41^ and have attempted to understand metabolic systems through modeling using genomic information from the community ^42,43^. Such genetic information may be important not only for improving our biological understanding of plant-microbe interactions but also for bioengineering applications. For example, DNA sequence motifs that are characteristic of important gene groups may be used as biomarkers, and SynComs can be further improved by functional modification via genetic engineering and/or by more efficient SynCom design.

The feature importance analysis of the model constructed in this study provides notable insight into interactions between organisms within the tomato rhizosphere. The most important features of the S-I model are microbial interactions, which suggests that the impact of inoculation with multiple microorganisms is significantly greater than the impact of inoculation with a single microorganism. In particular, the top two combinations—i.e., T2-2 with RC1 and T3-8 with T5-5—showed significantly higher growth when simultaneously inoculated (Fig. 5B). Subsequent multi-omics analyses suggested that RC1 was closely related to plant biomass, while T2-2 and T5-5 were microbial taxa that were characteristically dominant in the root microbiota following SynCom G2 inoculation (Fig. 9C). Although T5-5 was present in both SynCom G2 and B5, it was relatively more abundant in SynCom G2. This may be due to interactions between T5-5 and T3-8 in SynCom G2, which may in turn promote the colonization of T5-5 in the roots. Genome analysis revealed that both T2-2 and T5-5 possess genes involved in plant hormone metabolism (Fig. 2B). For example, the auxin biosynthesis gene possessed by T2-2 may be a representative PGP trait, since auxin-producing bacteria such as *Aeromonas punctata* and *Azospirillum brasilense* are known to stimulate plant growth and induce morphological alteration in Arabidopsis ^44–46^. Furthermore, the ACC deaminase gene possessed by T5-5 functions to degrade ethylene precursors and limit ethylene levels in plants exposed to stress conditions ^38^. For instance, ACC deaminase-producing bacteria such as *Bacillus* sp., *Brevibacterium linens*, and *Streptomyces venezuelae* have been found to enhance rice plant growth under salinity and heat stress ^47–50^. Considering these studies, T2-2 and T5-5 isolated here may play a central function in the promotion of tomato growth by SynCom G2.

Next we considered the interaction between microbes and metabolites, and the feature importance of the S-I model suggested that there was a crucial relationship between RC1 and tomatine. In a previous study from our team, we showed that tomatine, which is secreted by tomato roots into the rhizosphere, is involved in assembling the tomato rhizosphere microbiota and increases the abundance of *Sphingobium* spp. in the soil ^9^. More recently, we also found that RC1 has a tomatine degradation pathway ^10^. In this study, we found that tomatine enhanced the colonization of inoculum RC1 (Fig. 3A). This finding confirms these earlier reports, and the fact that simultaneous inoculation enhanced heat stress tolerance of tomato plants (Fig. 5C) suggested that the tomatine-mediated relationship between *Sphingobium* spp. and tomato is important for tomato stress tolerance. In contrast, such effects were not observed with the soybean-derived metabolite daidzein, implying that impact of plant-derived metabolites on the rhizosphere microbiota may be compound-specific. Multi-omics analysis also showed that among the many microbes present in the tomato rhizosphere, *Sphingobium* spp. are the most closely related to tomato growth (Fig. 10). Therefore, *Sphingobium* spp. may be a potential indicator of tomato growth. However, we also found that *Sphingobium* sp. RC1 did not have as many representative PGP trait-related genes as other SynCom members (Fig. 2B). Consequently, it may indirectly contribute as a helper microbe. For example, it may suppress the antimicrobial effect of tomatine, thereby promoting the colonization of other beneficial microbes and the development of a healthy microbial community within the rhizosphere. Moreover, the expression of Solyc05g053080 showed a strong positive correlation with both *Sphingobium* abundance and plant biomass (Fig. 10). Although its precise physiological function remains unclear, it may be involved in tomatine biosynthesis, potentially promoting tomato-*Sphingobium* interactions and contributing to enhanced plant growth.

Next, we note that characteristic changes in gene expression in tomatoes inoculated with SynCom G2 included an increase in the expression levels of HSF and HSP genes under high temperature conditions (Fig. 8) relative to the corresponding levels observed in non-inoculated and SynCom B5-inoculated tomatoes. This result indicates that SynCom G2 inoculation augmented the heat stress response of tomato plants, resulting in improved growth under heat stress conditions. Similar observations have been reported in other studies, where inoculations of ACC deaminase-producing *Brevibacterium* and thermotolerant *Bacillus* were found to increase HSP gene expression under heat stress and enhance the heat tolerance of rice and soybean, respectively. ^50,51^. In general, HSFs are key transcription factors regulating the heat stress response ^52^, while HSPs are molecular chaperones that prevent the aggregation of denaturing proteins and maintain protein homeostasis under heat and other forms of stress ^53^. In addition, in the plant biomass-associated network module (Fig. 10A), the hub gene Solyc05g053080 was found to be homologous to endomembrane-type CA-ATPase 4 (ECA4), a gene that has been found to be crucial for reactive oxygen species (ROS) production by RBOHD in *Arabidopsis thaliana* under salinity stress conditions ^54^. In general, RBOHD is critical for rapid systemic ROS signaling in response to diverse abiotic and biotic stimuli in plants ^55,56^. Moreover, ROS signaling leads to downstream HSF activation and HSP accumulation in the heat stress response ^57,58^. For the SynCom G2 inoculation designed in this study, it is possible that nonpathogenic or symbiotic bacterial stimuli may have activated these signaling pathways and thereby enhanced the heat stress response.

It should be noted that since this study analyzed gene expression only one month after SynCom inoculation, it is possible that some direct physiological responses to SynCom inoculation may not have been observed. Further detailed analysis of time-series changes after SynCom inoculation is expected to further elucidate the molecular mechanisms involved in SynCom colonization and plant response.

In conclusion, we demonstrate that a simple linear model can effectively predict and guide the design of synthetic microbial communities that enhance plant growth and stress resilience. We show that incorporating genomic information from SynCom members further improves prediction accuracy. In addition, we validate the effectiveness of model-designed SynComs in actual field cultivation. Finally, feature importance analysis of our model and multi-omics analysis suggests that specific microbial interactions within the rhizosphere microbial community play a key role in conferring abiotic stress tolerance in tomato plants.

## Materials and Methods

### Isolation and culturing of tomato endophytic bacteria

To collect root endophytes, bacterial strains were isolated from the roots of tomato plants (*S. lycopersicum* cv. Micro-Tom). Tomato seeds were sown in pots filled with a 1:1 mixture of vermiculite and field soil collected at the Kyoto University of Advanced Science, Kameoka, Kyoto, Japan (34°99’38”N, 135°55’14”E). Seedlings were then grown in the laboratory for seven weeks at 25°C under a 16h/8h light/dark cycle. At seven weeks we obtained samples of tomato roots, then carefully removed the soil adhering to the roots, including the rhizosphere and rhizoplane, following a previously described protocol ^59^. Next, endosphere compartments were washed with tap water before being homogenized in a 10 mM MgCl2 solution by mortar and pestle. The resulting homogenates were then diluted and distributed into agar plates containing six different media, i.e., Tryptic Soy Broth (TSB), Tryptone Yeast extract Glucose (TYG), Yeast Extract Mannitol (YEM), M408, M715, and Tap Water Yeast Extract (TWYE) media ^60^. All plates were incubated for 2–3 days at 28°C. Next, bacterial genomic DNA was extracted from each colony using the following method. First, colonies were suspended in 10 μL of an extraction buffer containing 25 mM NaOH and 0.2 mM EDTA heated at 95°C for 30 min. Next, 10 μL of 40 mM Tris-HCl solution (pH 6.8) was added. The *16S rRNA* genes in their respective isolates were then PCR-amplified using the following primer set: 10F (5′-GTTTGATCCTGGCTCA-3′) and 800R (5′-TACCAGGGTATCTAATCC-3′). PCR amplification used KOD FX Neo (TOYOBO, Osaka, Japan) and extracted DNA as the template sequence. The thermocycle program was as follows: 94°C for 2 min followed by 35 cycles of 98°C for 10 sec, 50°C for 30 sec, and 68°C for 1 min. The generated PCR products were then purified using a Wizard Genomic DNA Purification Kit (Promega, Madison, WI, USA) with all procedures performed as per the manufacturer’s instructions. After amplification, *16S rRNA* gene sequences were sequenced using the 10F primer, after which each bacterial isolate was annotated using BLAST search. All isolates were cultured on a growth medium (pH 7.0) containing 10 g L^−1^ peptone, 10 g L^−1^ beef extract, and 5 g L^−1^ NaCl, and were stored in 20% glycerol at −80°C. Isolates were passaged on agar plates containing the above growth medium.

### Genome analyses of SynCom member bacteria

Each bacterial member of a given SynCom was streaked onto a growth medium agar plate as described above. Next, a single colony was cultivated in 10–50 mL of liquid medium for three days at 28°C following preculture in 2 mL of the same medium. After incubation, these cultures were centrifuged at 4,000 × *g* for 5 min before being washed in 5 mL of TE buffer (i.e., 10 mM Tris-HCl (pH 8.0) and 1 mM EDTA). Cell pellets were then stored at −30°C for genomic DNA extraction. Genomic DNA of T2-2, T2-11, T2-21, and T5-5 was extracted using a Genomic DNA Buffer Set and a QIAGEN Genomic-tip 100/G column (QIAGEN, Hilden, Germany), with all procedures following the manufacturer’s instructions. Genomic DNA of T1-8, T2-14, T2-23, and T3-8 was extracted as per a previously described protocol ^61^ with minor modifications. Briefly, cell pellets were lysed with TE buffer containing lysozyme, sodium dodecyl sulfate, and proteinase K before being incubated at 37°C for 1 hour. Next, cleared lysate was pushed 10 times through a syringe (38 mm × 0.8 mm) before being deproteinized using TE-saturated phenol:chloroform:isoamyl alcohol (25:24:1). Genomic DNA was then collected via ethanol precipitation before being dissolved in 100 μL of TE buffer. DNA concentrations were measured using a BioSpec-nano (Shimadzu, Kyoto, Japan) and a Qubit 2.0 Fluorometer (Thermo Fisher Scientific, Waltham, MA, USA). After quantification, all genomic DNA samples were kept at 4°C until genome analysis.

Library construction and whole-genome sequencing were carried out as per a previously described protocol ^34^ with minor modifications. Briefly, a DNA library was cut off at 15 kbp (for T2-2, T2-11, T2-21, T2-23, and T5-5) or 10 kbp (for T1-8, T2-14, and T3-8) using a Blue Pippin size selection system (Sage Science, Beverly, MA, USA). Genomic libraries were then sequenced on a single PacBio sequel II system 2.0 cell. Using the resulting long-read data, genomes for T2-2, T2-11, T2-21, T2-23, and T5-5 were assembled using the Hierarchical Genome Assembly Process (HGAP) version 4 as implemented within SMRTlink version 10.0. Genomes of T1-8, T2-14, and T3-8 were assembled using the Improved Phased Assembler (IPA) implemented within SMRTlink version 10.2 according to their expected genome sizes. Circlator version 1.5.5 ^62^ was used to assess whether genome assemblies were circularizable and to predict the starting position. Next, genome-based species identification of SynCom members was performed using GTDB-Tk version 2.1.0 using the default parameters ^63^. Gene prediction within genomes was performed using Prokka version 1.14.6 ^64^. The functional annotation of predicted gene sets was performed using eggNOG-mapper version 2.1.9 using the default parameters ^65,66^. For plant growth-promoting traits, pathways related to auxin production, suppression of ethylene accumulation, phosphate solubilization, siderophore production, and nitrogen fixation were retrieved from the MetaCyc ^67^ and KEGG pathway databases ^68^. The names and KEGG Orthology IDs of genes involved in these pathways were queried in the functional annotations of the respective genome. Using this process, we created a plant growth-promoting (PGP)-related gene possession profile from combined sets of genes. Finally, orthologous gene groups of SynCom member genomes were inferred using OrthoFinder version 2.5.4 using the default parameters ^69^.

### Laboratory growth experiments of SynCom-inoculated tomato plants

We first sterilized tomato seeds before sowing them on Murashige–Skoog medium supplemented with 3% sucrose and 0.3% gellan gum as per a previously described protocol ^9^. Next tomato seedlings were grown at 25°C for two weeks before they were subjected to SynCom inoculation. For each SynCom preparation, a particular configuration was precultured in 2–3 mL liquid growth medium before being cultivated in 75–500 mL of the same medium for three days at 28°C. Cultures were then centrifuged at 4,000 × *g* for 5 min before being washed with 20 mL of autoclave-sterilized tap water. Cell pellets were then resuspended in 20 mL sterile tap water. Next, two-week-old tomato seedlings were transferred to pots containing 40 g vermiculite. The inoculation mixture contained 100mL sterile tap water containing 0.2% (w/v) Otsuka House No. 1 and No. 2 standard nutrients (Otsuka Chemical Co., Osaka, Japan) as well as a cell suspension of each SynCom member at OD600 = 0.05; these solutions were inoculated into tomato seedlings. Plants grew at 25°C for 14 days before being assigned one of three different growth temperatures (i.e., 30°C, 35°C, or 38°C) in which they grew for another two weeks. Throughout the cultivation period all tomato plants were grown under a 16h/8h light/dark cycle. After cultivation, tomato plants were separated into aerial parts and roots. The roots were rinsed with tap water, and any residual water was removed using Kimwipes. Finally, we measured the fresh weight of the above ground parts and roots of the tomato plants.

### Microbiome analysis of endophytic bacteria in tomato roots

After fresh weight measurements were taken, SynCom-inoculated tomato roots were homogenized using a BioMasher® Ⅱ (Nippi. Inc., Tokyo, Japan). Next, DNA was extracted from the resulting homogenate using a DNeasy PowerSoil Pro Kit (QIAGEN) with all procedures performed as per the manufacturer’s instructions. Next, amplicons of the 16S rRNA V4 region were prepared from each root DNA sample as per a previously described protocol ^9^. In brief, the V4 region was PCR-amplified using a 515F/806R primer set with mitochondrial- and chloroplast-specific peptide nucleic acids (mPNA [N-term-GGCAAGTGTTCTTCGGA-C-term] and pPNA [N-term-GGCTCAACCCTGGACAG-C-term], respectively [PANAGENE Inc, Daejeon, Republic of Korea]). This was done to prevent the amplification of tomato mitochondrial and plastid sequences ^70^. After PCR, the resulting amplicons were purified using an AMPure XP kit (Beckman Colter, Danvers, MA, USA) before being attached to a MiSeq (Illumina, San Diego, CA, USA) adapter via PCR. After purification of PCR products, the prepared 16S amplicon mixture was sequenced on a MiSeq (Illumina) platform by FASMAC Co. Ltd to obtain 2 × 250-bp paired-end sequences.

Raw sequences were processed using the QIIME2 version 2021.2 pipeline ^71^. Raw paired-end sequences were first trimmed before the first 20 bases and after 200 bases. Next, denoised amplicon sequence variants (ASVs) were generated using DADA2 ^72^ using the q2-dada2 plugin in QIIME2. Taxonomic classification of ASVs was performed using the q2-feature-classifier plugin and a Naive Bayes classifier ^73^, which has been pretrained on operational taxonomic units (99% identity) from the 515F/806R region of sequences in the SILVA rRNA gene database release 138 ^74^. After filtering out mitochondrial and chloroplastic sequences, we obtained 20,141–374,296 reads per sample, representing 8,603 ASVs, and these were rarefied to 20,000 reads per sample for subsequent diversity analysis. Next, alpha and beta diversity indices were computed using the core-metrics-phylogenetic pipeline of the q2-diversity plugin. This plugin also performed principal coordinate analysis (PCoA) using weighted and unweighted UniFrac distances. Multiple alignment of ASVs was performed using MAFFT ^75^ and a phylogenetic tree was generated by FastTree ^76^, both implemented within the q2-phylogeny plugin. Finally, metagenomic functional abundance profiles were estimated using PICRUSt2 ^77^ and the KEGG pathway database.

Microbiome analysis of SynCom G2- and SynCom B5-inoculated tomato plants was performed using the same sequencing protocol described above. Here, we obtained 66,497– 103,566 reads per sample representing 1,290 ASVs. These were then rarefied to 66,000 reads per sample for subsequent diversity analysis.

All statistical analyses of microbiome datasets were performed using R. The R package *stats* was used for Wilcoxon tests, the package *vegan* for PERMANOVA tests ^78^, and *ALDEx2* for differential abundance analysis ^79,80^.

### Model construction

As the response variable, plant shoot biomass values were first converted to ratios with control plants (i.e., noninoculated, 30°C) for each cultivation trial. These values were then box-cox transformed. For all models, the 301 samples were split 7:3 between training and test datasets for model construction and subsequent performance evaluation. This step was randomly repeated 50 times to evaluate average performance relative to variation in the training and test data split. Also, the hyperparameters of each model were determined using the five-fold cross-validation method.

For SynCom-Inoculant models, the presence or absence of microbes and metabolites in each SynCom was expressed as 1/0 and the cultivation temperatures (i.e., 30°C, 35°C, and 38°C) were scaled from 0 to 1. These variables and the interaction terms between each variable pair were used as the set of explanatory variables. For the SynCom-Genome model, the gene composition of each SynCom was estimated based on orthologous gene possession profile of constituent SynCom bacteria, and the presence or absence of orthologous gene groups was expressed as 1/0. These variables, the presence or absence of metabolites, and scaled cultivation temperature values were used as explanatory variables. For the Microbiome model, taxonomy profiles at the Phylum and Family levels and KEGG pathway profiles were filtered to the taxon and pathway detected in at least 5% (15 samples) of the 301 samples. The filtered taxon and pathway count data were centered log-ratio (CLR) transformed. These variables, the presence or absence of metabolites, and scaled cultivation temperature values were used as explanatory variables.

The following R packages: *ROCR*, *caret*, *glmnet*, *lme4*, and *randomForest* were used for machine learning ^81–84^.

### Validation of SynCom model-based design

The S-I model was used to predict plant growth for all SynCom compositions and growing temperatures (i.e., 30°C, 35°C, and 38°C). We selected those SynCom compositions predicted to be in the top 5% of predicted growth outcomes for each growing temperature, classified them into eight groups using the k-means method, and then selected one from each group to validate the eight unexplored SynCom compositions. We used a similar procedure to identify “bottom 5%” and “10% around the median” SynCom compositions. These were also classified into eight groups and a representative was then selected, yielding a total of 24 newly designed SynCom compositions. Laboratory growth experiments were conducted as described above.

### Field trials of SynCom-inoculated tomato plants

Tomato seeds (*S. lycopersicum* cv. Aiko) were sown on commercial soil (Nihonhiryo com.) and grown for 2 weeks in a growth chamber (25°C, 16h/8h light/dark photoperiod). On the day of inoculation, seedlings were transplanted to pots (diameter 9 cm, height 7 cm) filled with 200 g of soil containing a tomatine aqueous solution (25 nmol/g soil); specifically, we added 80 ml tap water containing 5 μmol tomatine (Tokyo Chemical Industry) to each measured quantity of 200 g soil. Each pot was then placed in a plastic bag to prevent water loss. Next, prepared (see above) SynCom bacterial suspension solutions were inoculated onto the seedlings. The inoculation was performed such that 12.5 μL aqueous solution of each SynCom (OD600 = 0.5) was included in 1 gram of soil. Seedlings were then grown for 2 weeks of acclimatization in the greenhouse in early summer (i.e., June 24–July 8, 2022) before being moved to a field site at the Tokyo University of Agriculture and Technology, Fuchu, Tokyo, Japan (35°68’54”N, 139°48’04”E). Each plot was one square meter in size, and contained four plants of the same treatment. Plots were separated by 50 cm of empty space, and four plots were created for each SynCom group, including the control. In total, 96 plants were cultivated for one month in the field. Fresh weights of aerial/aboveground and root portions of plants were measured immediately after sampling. Plant dry weight was then estimated after oven-drying at 80°C until reaching a constant weight.

### Transcriptomic analysis of SynCom-inoculated tomato plants

Fully-expanded upper leaves and roots were sampled from SynCom-inoculated tomato plants, then were ground in liquid nitrogen using a mortar and pestle. Total RNA was extracted using RNeasy Plant Mini Kits (QIAGEN) and a RNase-Free DNase Set (QIAGEN), with all procedures following the manufacturer’s protocol. Next, RNA-seq libraries were constructed using the Lasy-Seq version 1.1 method ^85^. Finally, prepared libraries were sequenced on a HiSeqX (Illumina) platform to obtain 2 × 150-bp paired-end read datasets.

Raw reads were first subjected to quality control by removing low-quality bases using Trimmomatic ^86^ using the default parameters. Trimmed reads were then aligned to the tomato genome (Solanum_lycopersicum. SL3.0) using STAR version 2.7.6a ^87^ and the Ensembl Plants release 50 ^88^ gene annotation framework. Gene expression levels were estimated as transcripts per million (TPM) using RSEM version 1.3.3 ^89^ using the default parameters.

All statistical analyses were performed using R. Of the 25,366 genes detected, 14,948 genes in the leaf and 16,727 genes in the root were kept for downstream analysis following the exclusion of low expression genes (i.e., mean TPM < 1). The *DESeq2* R package was used for differential expression analysis ^90^. GO annotation of tomato genes was obtained from Ensembl Plants using the *biomaRt* R package ^91^, and enrichment analysis was performed using the *GOstats* and *GSEABase* R packages ^92^. Finally, gene expression patterns were clustered and visualized using the *pheatmap* R package.

### Integrated multiomics analyses

Sparse multiple canonical correlation analysis was performed on plant growth (i.e., shoot and root biomass) data, gene expression profiles (i.e., TPM value, including 14,948 genes from leaves and 16,727 genes from roots), and root endophytic bacteria profiles (CLR value, 361 taxa at genus level), using the *PMA* R package ^93^. Parameter tuning was performed using 100 permutations and five iterations. Based on the sparse canonical variates of the first and second components, we extracted 4,938 genes in the leaves and 7,089 genes in the roots, representing 176 taxa.

Next, a multi-omics co-occurrence network was created using WGCNA ^94^ using a previously described protocol with some modification ^59^. Briefly, a signed network was created using the Spearman correlation matrix as well as the soft thresholding power selected from a scale-free topology fit index R-squared value of 0.75. Module identification was then performed using a dynamic tree cut based on a hierarchical clustering tree. Network connections with a topological overlap above a threshold of 0.08 were extracted, and the resulting subnetwork was visualized using the *igraph* R package ^95^.

## Data availability

The genome assemblies of SynCom member bacteria were deposited in DDBJ/ENA/GenBank under accession numbers AP027908–AP027921. Raw sequence data for genome analysis were deposited at the DDBJ Sequence Read Archive (DRA) under accession numbers DRR453250–DRR453257 (BioProject accession number PRJDB15212). Raw data describing SynCom composition and plant growth used for modeling are provided in Supplemental Table S3.

## Acknowledgments

This work was supported by JST CREST (Grant Number JPMJCR15O2 and JPMJCR17O2) and GteX Program (JPMJGX23B2), and RIKEN TRIP initiative to K.S., Japan.

