## Supplemental Figures for "From model to field: predictive design of microbial communities for resilient crop performance"

**Title**

### Supplemental Figures

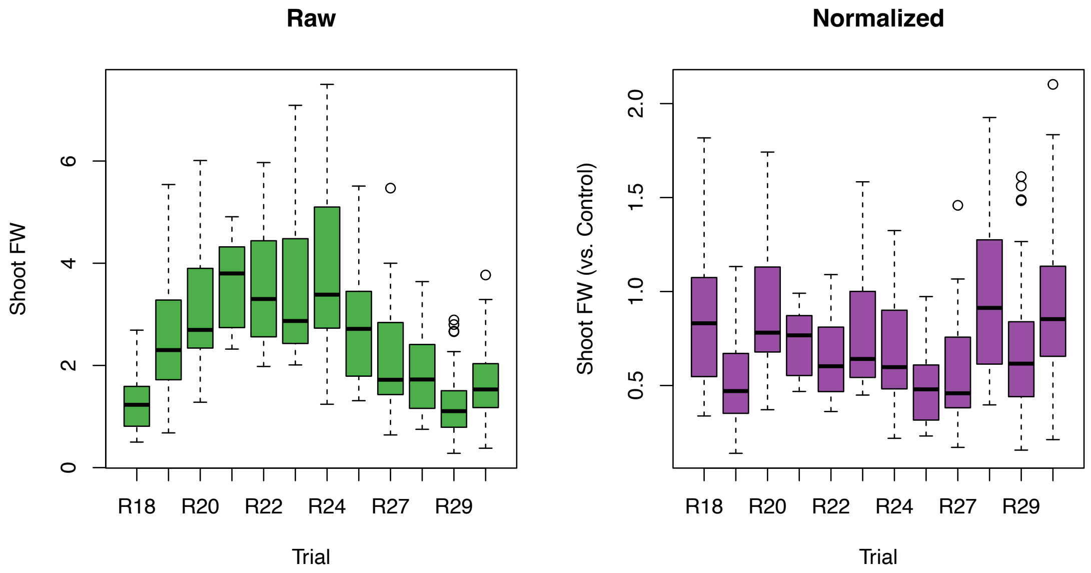


Supplemental Fig. S1. Plant growth in the series of trials. Raw data of plant biomass (shoot fresh weight) and batch corrected data with the control conditions.

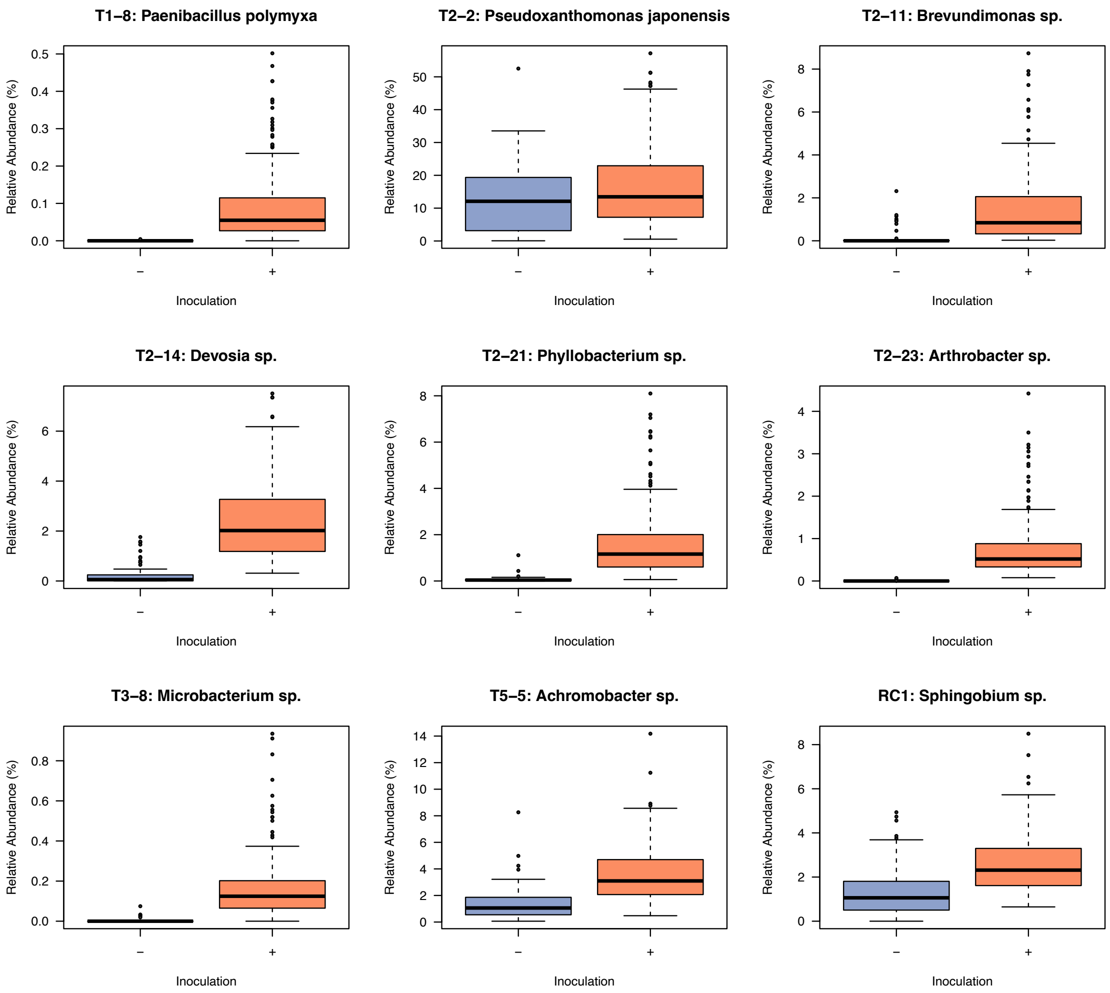


Supplemental Fig. S2. Relative abundance of ASVs identical to each inoculant +/− inoculation.

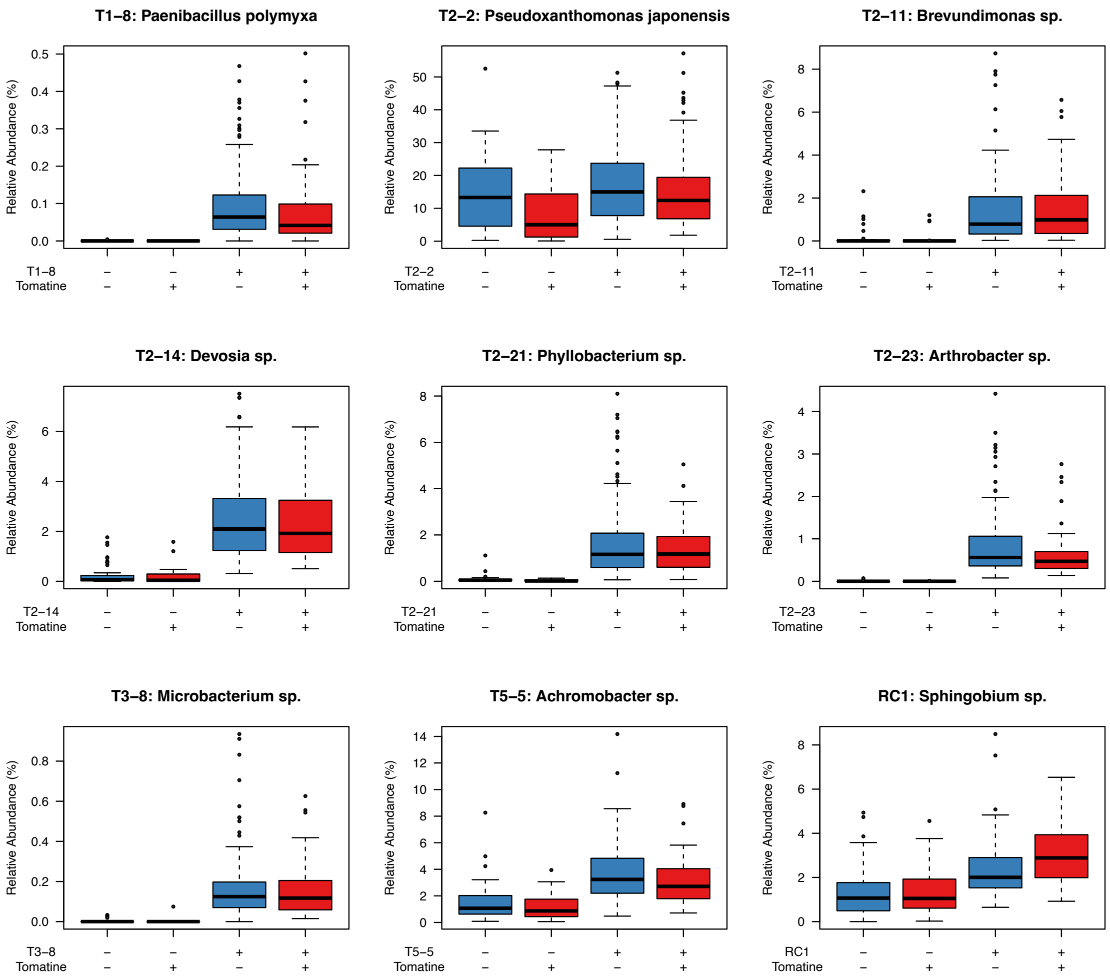


Supplemental Fig. S3. Relative abundance of ASVs identical to each inoculant +/− inoculation with tomatine.

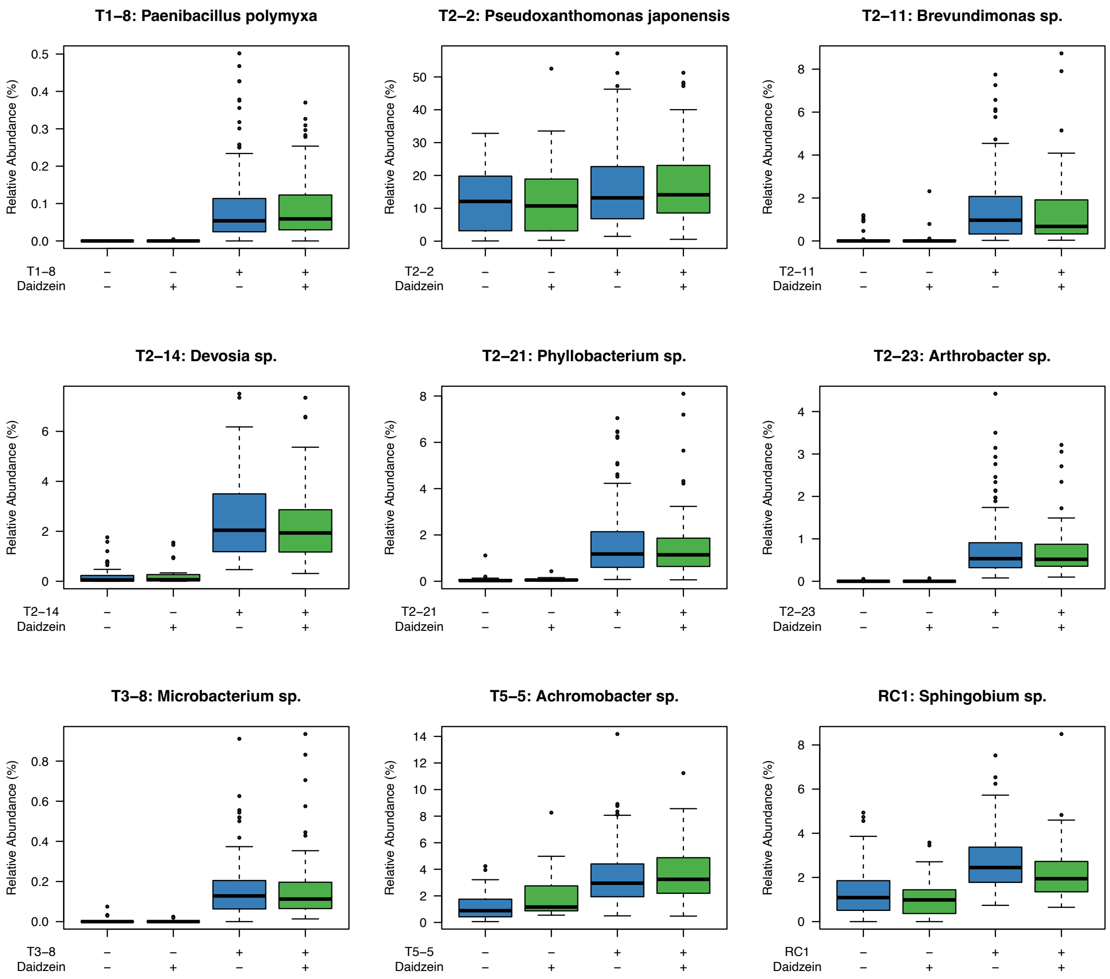


Supplemental Fig. S4. Relative abundance of ASVs identical to each inoculant +/− inoculation with daidzein.

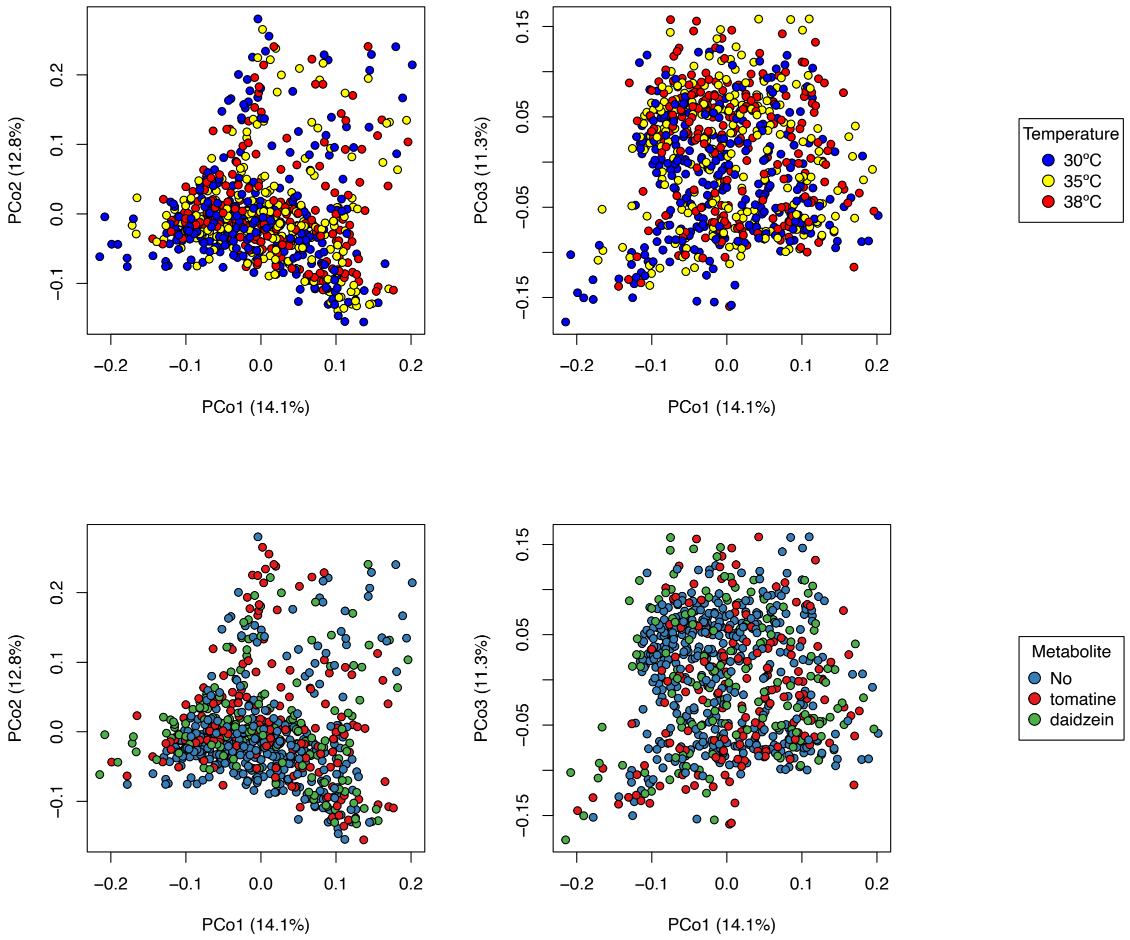


Supplemental Fig. S5. Principal coordinate analysis plot based on weighted UniFrac distance of microbiome data for modeling.

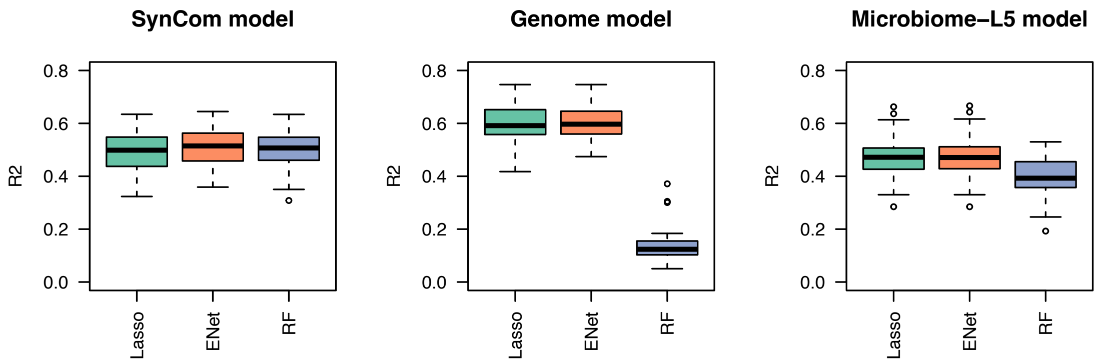


Supplemental Fig. S6. Comparison of prediction performance among different model algorithms.

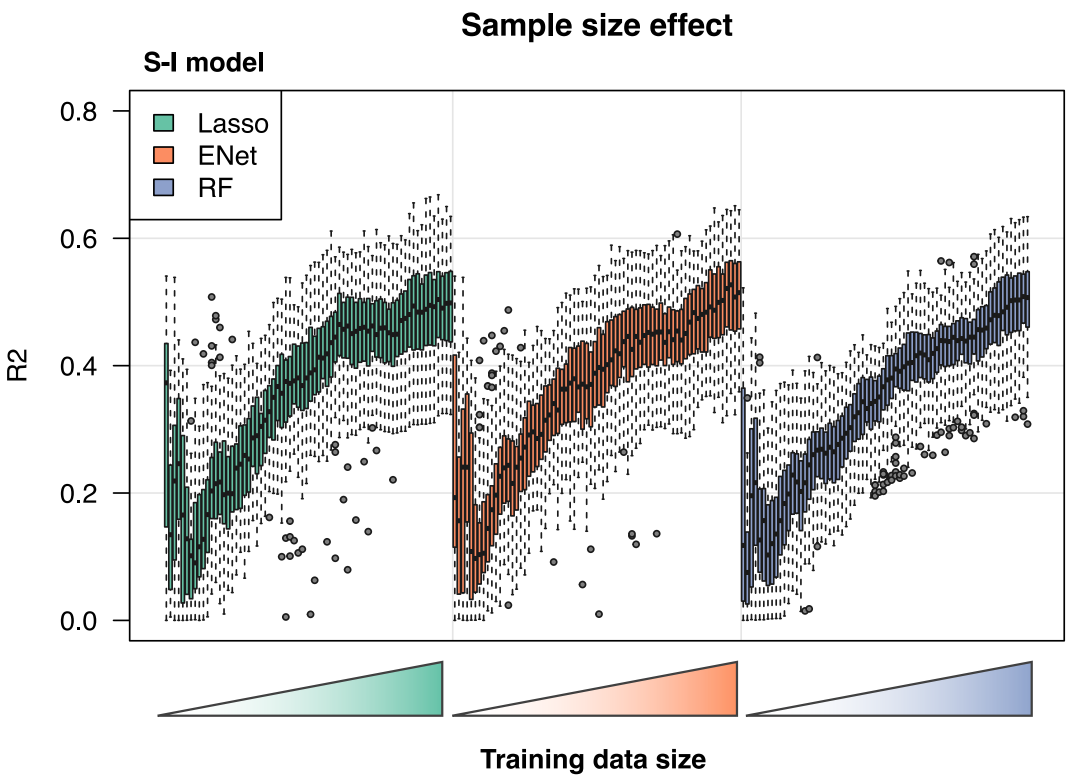


Supplemental Fig. S7. Prediction performance affected by the size of training data set for modeling.

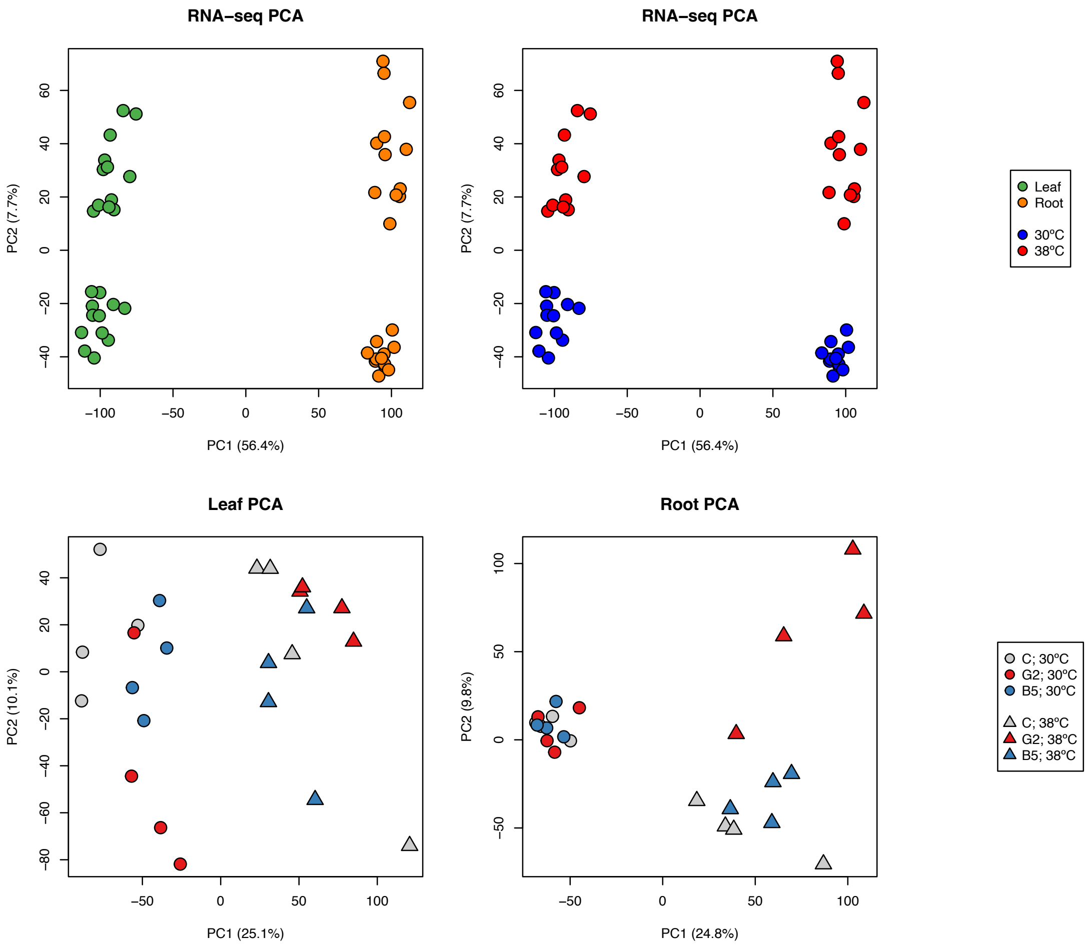


Supplemental Figs. S8. Principal component analysis plots of transcriptome data.

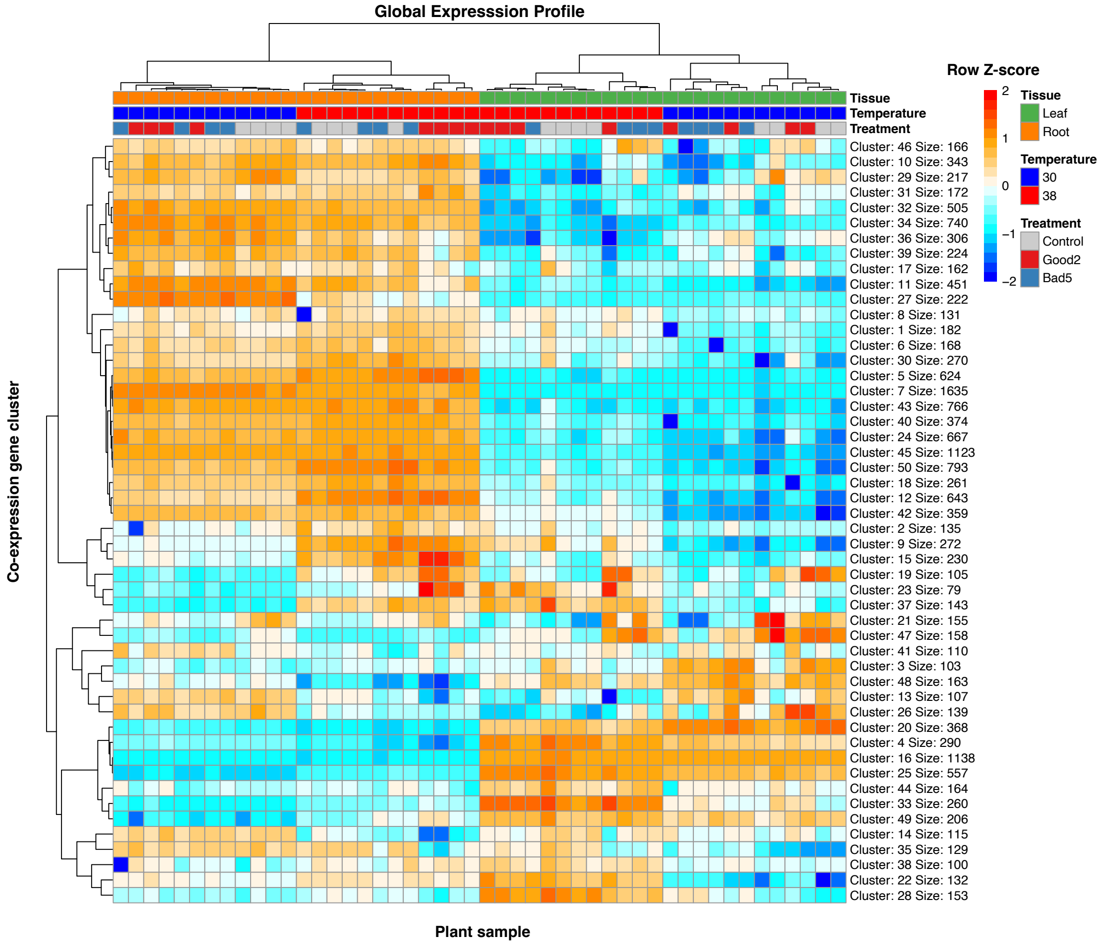


Supplemental Figs. S9. Heatmap of gene expression profile. Genes were clustered by k-means method. The optimal k was determined based on the Bayesian information criterion. Representative expression patterns of genes in each cluster were shown by heatmap.

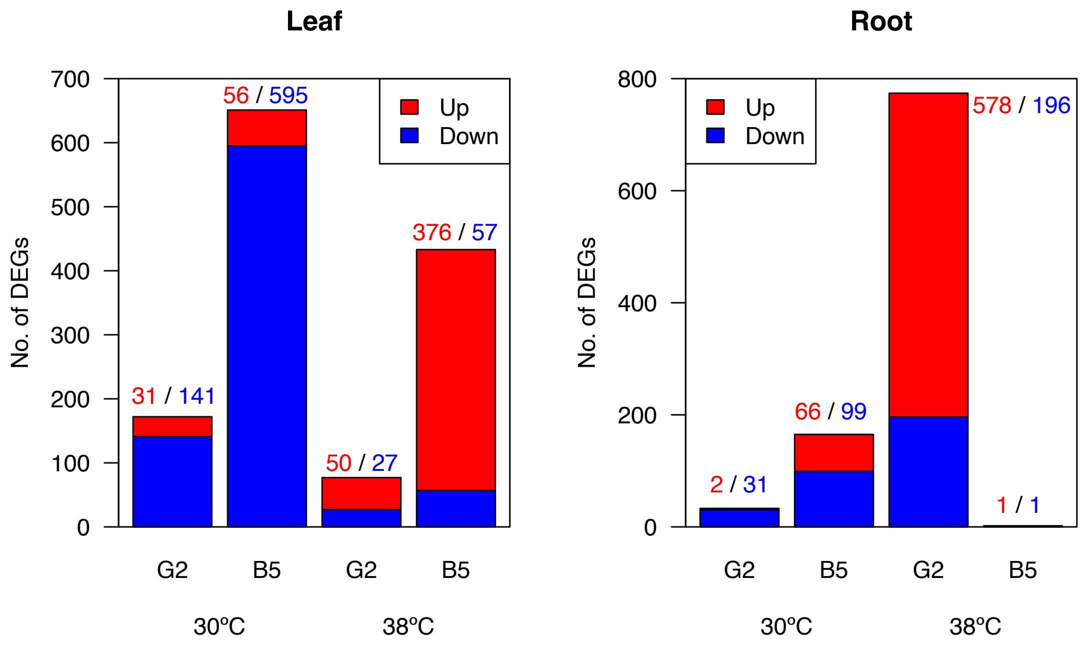


Supplemental Fig. S10. Number of differentially expressed genes by SynCom inoculation. Differentially expressed genes by SynCom G2 and B5 inoculations were identified for each tissue and temperature (FDR < 0.01, DESeq2).

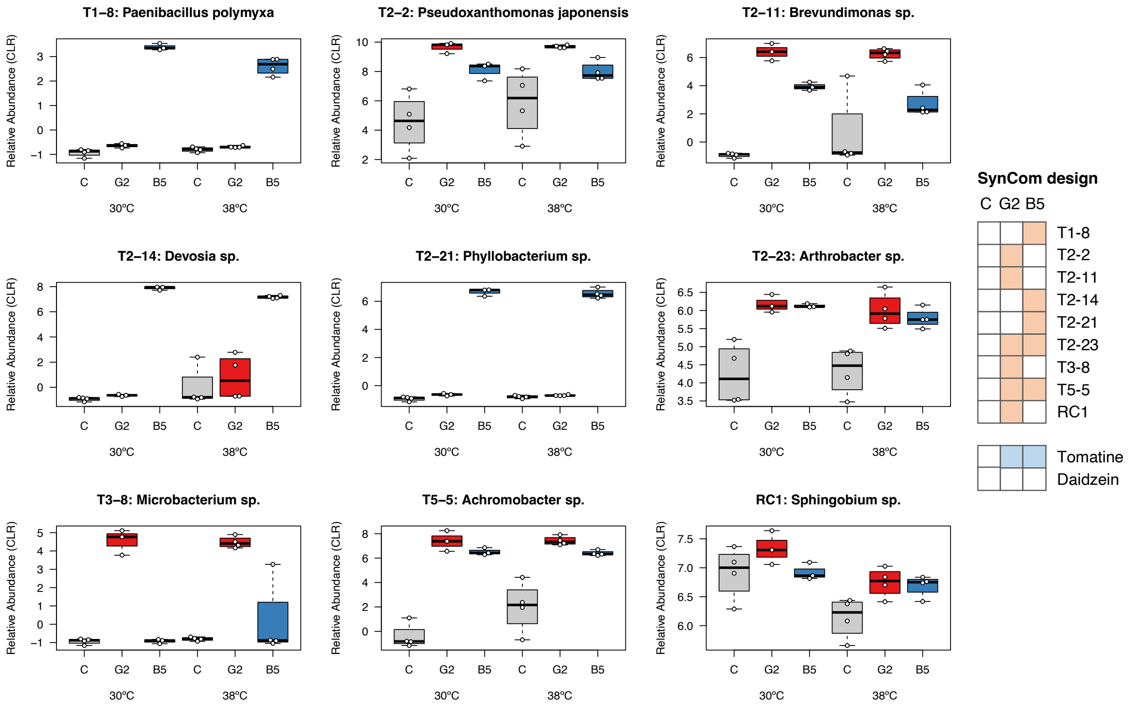


Supplemental Fig. S11. Relative abundance of ASVs identical to each inoculant +/− SynCom inoculation.
